# SirA inhibits the essential DnaA:DnaD interaction to block helicase recruitment during *Bacillus subtilis* sporulation

**DOI:** 10.1101/2022.04.18.488658

**Authors:** Charles Winterhalter, Daniel Stevens, Stepan Fenyk, Simone Pelliciari, Elie Marchand, Panos Soultanas, Aravindan Ilangovan, Heath Murray

## Abstract

Bidirectional DNA replication from a chromosome origin requires the asymmetric loading of two helicases, one for each replisome. Our understanding of the molecular mechanisms underpinning helicase loading at bacterial chromosome origins is incomplete. Here we report both positive and negative mechanisms for directing helicase recruitment in the model organism *Bacillus subtilis*. Systematic characterization of the essential initiation protein DnaD revealed distinct protein interfaces required for homo-oligomerization, interaction with the master initiator protein DnaA, and interaction with the helicase co-loader protein DnaB. Informed by these properties of DnaD, we went on to find that the developmentally expressed repressor of DNA replication initiation, SirA, blocks the interaction between DnaD with DnaA, thereby inhibiting helicase recruitment to the origin during sporulation. These results advance our understanding of the mechanisms underpinning DNA replication initiation in *B. subtilis*, as well as guiding the search for essential cellular activities to target for antimicrobial drug design.

## INTRODUCTION

Genome replication initiates at specific chromosomal loci termed origins. Throughout the domains of life, initiator proteins containing a conserved AAA+ (*A*TPase *A*ssociated with various cellular *A*ctivities) motif assemble at chromosome origins and direct loading of two helicases for the onset of DNA replication (1). Interestingly, while the initiation pathway in both bacteria and eukaryotes culminates in a ring shaped hexameric helicases encircling a single DNA strand, the molecular mechanisms necessary to achieve this crucial helicase loading step appear to be distinct (2). Bacteria use their master initiator DnaA to first unwind the chromosome origin (*oriC*) and then load helicases around ssDNA such that they are poised to start unwinding. The eukaryotic initiator ORC (*O*rigin *R*ecognition *C*omplex) also promotes helicase loading, but in this case the annular enzyme is deposited around double-stranded DNA (dsDNA) in a dormant state which must subsequently be activated to form an open complex and encircle a single strand. These distinctions make bacterial DNA replication initiation proteins attractive targets for the development of novel antibiotics (3–5).

Despite decades of study, the molecular mechanisms coordinating helicase recruitment and loading during the bacterial cell cycle are unclear (6–9). Moreover, bacteria are not known to directly regulate helicase recruitment, instead they are thought to regulate the ability of the ubiquitous master initiator DnaA to bind and unwind the chromosome origin, ultimately initiating DNA replication in bacteria {Leonard, 2015 #2821}{Katayama, 2010 #2425}.

DnaA is a multifunctional enzyme composed of four distinct domains that act in concert during DNA replication initiation (Fig. S1A) (10). Domain IV contains a helix-turn-helix dsDNA binding motif and a basic loop with a conserved arginine that specifically recognize 9 base-pair asymmetric sequences called “DnaA-boxes” (consensus 5′- TTATCCACA-3′) (11–13). Domain III is composed of the AAA+ motif that can assemble into an ATP-dependent right-handed helical oligomer (14–16). Domain III also contains residues required for a DnaA oligomer to interact specifically with a trinucleotide ssDNA binding element termed the “DnaA-trio” (consensus 3′-GAT-5′) (17–19). It has been proposed that a DnaA oligomer, guided by DnaA-boxes and DnaA-trios within *oriC*, interacts with one strand of the DNA duplex to promote chromosome origin opening (17,19–21).

DnaA domain II tethers domains III/IV to domain I, which acts as an interaction hub. Domain I (DnaA^DI^) facilitates homo-oligomerisation, either directly through self-interaction (22) or indirectly via accessory proteins such as DiaA and HobA (23, 24). Domain I also interacts with important regulatory proteins such as HU, Dps and SirA (25–28) and has weak affinity for ssDNA (29). However, the most important role of DnaA^DI^ is thought to be recruiting the replicative helicase. This may occur either directly, as for *Escherichia coli* DnaA (30), or indirectly, as for *B. subtilis* DnaA (Fig. 1A) (31, 32). Interestingly, in both cases the same surface of DnaA domain I is suggested to be involved (Fig. S1B-C) (29,31,33–35).

**Figure 1.**
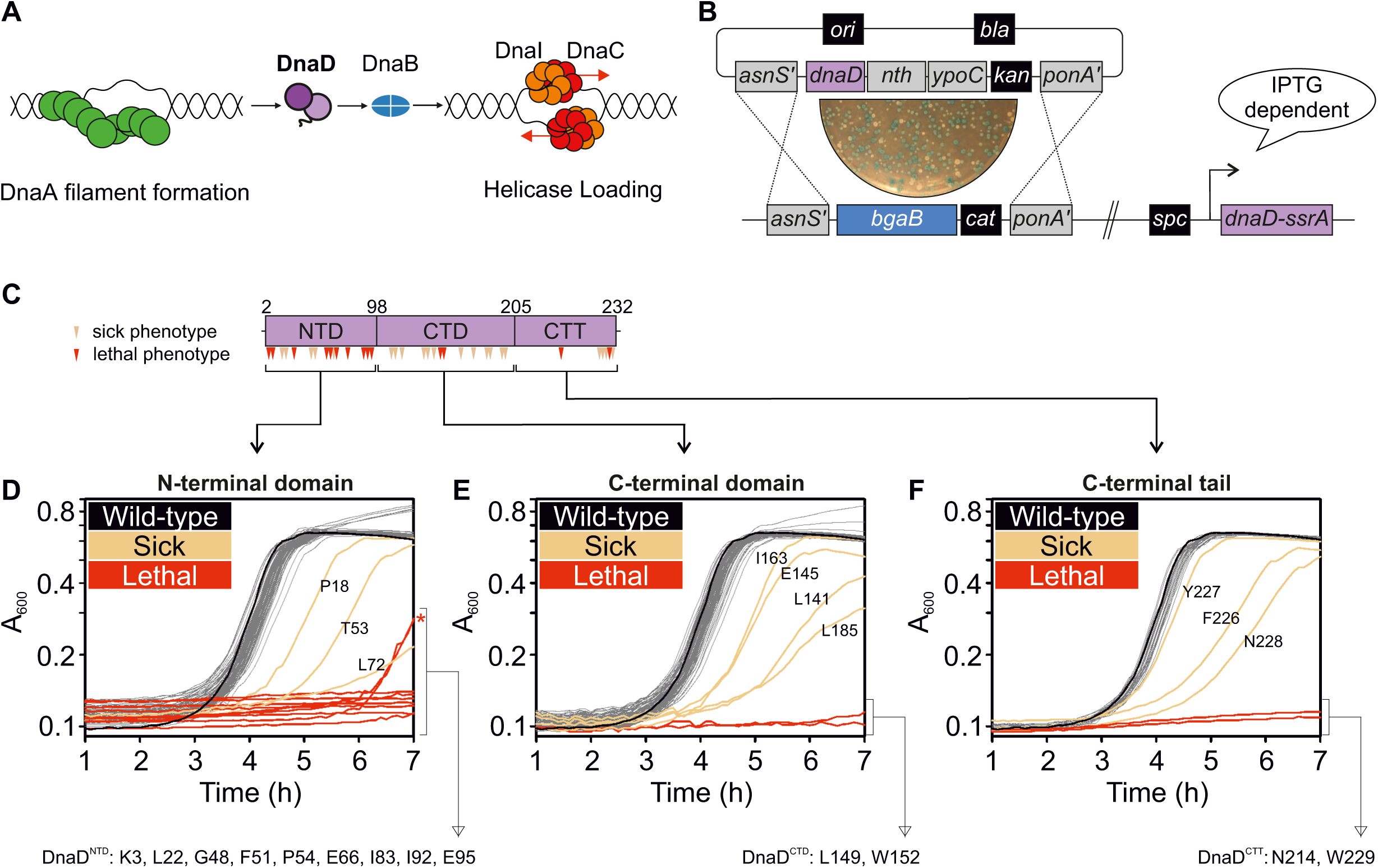
Identification of essential residues in *B. subtilis* DnaD. **(A)** Schematics of the helicase loading pathway in *B. subtilis* showing sequential recruitment of DnaA, DnaD, DnaB and the helicase complex DnaI-DnaC. **(B)** DnaD blue/white screening assay. An integration vector carrying individual *dnaD* substitutions is integrated by double recombination at the *dnaD* locus. Plasmid variants were constructed from pCW66. The recipient strain (CW197) is kept alive by the induction of the ectopic *dnaD-ssrA* with IPTG and selection of white colonies following transformation of *dnaD* substitutions is performed in the presence of kanamycin, the chromogenic substrate X-Gal and IPTG. **(C)** DnaD primary structure mapped with substitutions that conferred a different phenotype than wild-type. Residues are marked as red for lethal and beige for altered growth (translucent/heterogeneous phenotype or slow growth). NTD denote the N-Terminal Domain, CTD the C-Terminal Domain and CTT the C-Terminal Tail of DnaD. **(D-F)** Plate reader growth assays of DnaD variants located within the protein N-terminal domain **(D)**, C-terminal domain **(E)** or C-terminal tail **(F)** in the absence of DnaD-SsrA. The black line shows the growth profile of cells harboring wild-type *dnaD*, whereas beige lines highlight slow-growing variants and red line lethal substitutions. Note that some lethal substitutions develop suppressor mutations that activate the expression of *dnaD-ssrA* (red star). Lists of the DnaD variants that revealed a lethal phenotype by growth in plate reader assays are provided below panels (D-E): K3 (CW289), L22 (CW292), G48 (CW293), F51 (CW174), P54 (CW295), E66 (CW311), I83 (CW170), I92 (CW297), E95 (CW166), L149 (CW308), W152 (CW302), N214 (CW283), W229 (CW286). All strains in (D-E) are referenced in Table S6, the wild-type control is CW162 and the parent strain of all DnaD mutants is CW197.

Along with DnaA, DnaD and DnaB are required for the recruitment of the DnaC:DnaI complex (helicase and AAA+ chaperone, respectively) to the *B. subtilis* chromosome origin. Recruitment of initiation proteins to *oriC* has been shown to follow a linear pathway (Fig. 1A) (32,36,37). While DnaD and DnaB are known to be essential factors during both replication initiation and restart at repaired replication forks (38), a mechanistic understanding of the activities performed by these replication proteins has remained elusive (31,32,39,40).

In this paper we focus on DnaD. DnaD has three functional regions: N-terminal domain (DnaD^NTD^), C-terminal domain (DnaD^CTD^) and C-terminal tail (DnaD^CTT^) (Fig. 1C). The crystal structure of DnaD^NTD^ revealed a homodimer with a large interface stabilizing the two monomers (41). Also the DnaD^NTD^ was observed to be a tetramer in solution (42), hence a model of the DnaD dimer:dimer interface was proposed based on the observed symmetry of DnaD^NTD^ dimers packed within the crystal (41). Furthermore it has been suggested that the prominent winged-helix motif within the DnaD^NTD^ might act as a scaffold for the formation of higher order species (36). However, the DnaD^CTD^ structure was solved as a monomer {Marston, 2010 #2909} and the functional significance of DnaD oligomerization *in vivo* was unknown.

The DnaD activities required for *B. subtilis* DNA replication initiation have yet to be clearly defined. An interaction with DnaA was proposed to involve residues located in both the DnaD^NTD^ and the DnaD^CTD^ (31, 34). An interaction with DnaB was suggested to involve the DnaD^NTD^, and the DnaD^NTD^ and DnaB^NTD^ are predicted to be structurally related (31, 37). Again, the functional significance of specific DnaD interactions with DnaA and DnaB *in vivo* is unknown.

To explore the role of *B. subtilis* DnaD in the mechanism of DNA replication initiation, we performed a systematic alanine scan to identify residues essential for DnaD activities within its physiological environment of a cell. Structural and functional characterization of DnaD identified residues required for homo-oligomerisation and for protein:protein interactions with DnaA and DnaB. Guided by these discoveries, we went on to find that the developmentally expressed regulator of DNA replication initiation, SirA, blocks the interaction of DnaA with DnaD, thus inhibiting helicase recruitment during the early stages of sporulation to impede further rounds of DNA replication. Conservation of DnaD in several bacterial pathogens suggests that the DnaA:DnaD interface is an attractive target for rational drug design.

## MATERIALS AND METHODS

### Reagents

Nutrient agar (NA; Oxoid) was used for routine selection and maintenance of both *B. subtilis* and *E. coli* strains. Supplements were added as required: ampicillin (100 µg/ml), chloramphenicol (5 µg/ml), kanamycin (5 µg/ml), spectinomycin (50 µg/ml in *B. subtilis*, 100 µg/ml in *E. coli*), tetracycline (10 µg/ml), erythromycin (1 µg/ml) in conjunction with lincomycin (25 µg/ml), X-gal (0.01% v/v), xylose (0.35% v/v), IPTG (0.1 mM unless indicated otherwise). All chemicals and reagents were obtained from Sigma-Aldrich unless otherwise noted. Antibodies were purchased from Eurogentec. Plasmid extractions were performed using Qiagen miniprep kits. Other reagents used for specific techniques are listed within the method details.

### Biological resources: B. subtilis strains

*B. subtilis* strains are listed in Table S1 and were propagated at 37°C in Luria-Bertani (LB) medium unless stated otherwise in method details. Transformation of competent *B. subtilis* cells was performed using an optimized two-step starvation procedure as previously described (43, 44). Briefly, recipient strains were grown overnight at 37°C in transformation medium (Spizizen salts supplemented with 1 μg/ml Fe-NH_4_-citrate, 6 mM MgSO_4_, 0.5% w/v glucose, 0.02 mg/ml tryptophan and 0.02% w/v casein hydrolysate) supplemented with IPTG where required. Overnight cultures were diluted 1:17 into fresh transformation medium supplemented with IPTG where required and grown at 37°C for 3 hours with continual shaking. An equal volume of prewarmed starvation medium (Spizizen salts supplemented with 6 mM MgSO_4_ and 0.5% w/v glucose) was added and the culture was incubated at 37°C for 2 hours with continuous shaking. DNA was added to 350 μl cells and the mixture was incubated at 37°C for 1 hour with continual shaking. 20-200 μl of each transformation was plated onto selective media supplemented with IPTG where required and incubated at 37°C for 24-48 hours. The genotype of all chromosomal *dnaA* and *dnaD* mutants was confirmed by DNA sequencing. Descriptions, where necessary, are provided below.

CW197 [*trpC2* Δ*dnaD amyE*::*spc*(*P_HSA+1T_-dnaD-ssrA lacI^Q18M/W220F^)*] was constructed to study potentially lethal mutants of *dnaD in vivo* (Fig. 1B). First, an ectopic copy of *dnaD* was placed under the control of an IPTG-inducible promoter (*P_HYPERSPANK_*) (8). When combined with a deletion mutant of the native *dnaD*, the basal expression level of the ectopic copy was sufficient to sustain growth. Two approaches were taken to reduce the basal expression of the ectopic *dnaD*, lowering promoter activity and reducing DnaD stability. Promoter activity was inhibited by altering the transcription start site from A to T (*P_HAT_*) and introducing mutations into *lacI* (Q18M/W220F) that increase operator binding (Fig. S2A) (45, 46). DnaD stability was reduced by fusing the ectopic *dnaD* to an *ssrA* degradation tag (AANDENYSENYALGG) (47). Together these modifications produced a suitable expression system that conditionally complements the *dnaD* deletion mutant only when the ectopic *dnaD-ssrA* is induced (Fig. S2B). Immunoblot analysis confirmed nearly complete degradation of DnaD-ssrA following removal of IPTG within 30 minutes (Fig. S2C).

CW252 [*trpC2 amyE::spec(lacI P_HYPERSPANK_-sirA) ganA::erm(xylR P_XYL_-dnaD)*] was constructed by transformation with a PCR product generated by three-way Gibson assembly (NEBuilder HiFi). The *dnaD* gene with its native ribosome binding site was amplified using oCW611 and oCW612 using 168CA genomic DNA as template. The flanking region containing *ganA*′*-xylR-P_XYL_* was amplified using oCW284 and oCW614 with pJMP1 as template. The flanking region containing *erm-*′*ganA* was amplified using oCW613 and oCW130 with pJMP1 as template.

CW270/279/280 [*trpC2 amyE::spec(lacI P_HYPERSPANK_-sirA) ganA::erm(xylR P_XYL_-dnaD*)* with * being I83A, F51A and E95A were constructed identically to CW252 with the exception of *dnaD^F51A^, dnaD^I83A^* and *dnaD^E95A^* being amplified using oCW611 and oCW612 from CW174, CW170 and CW166 genomic DNA, respectively as templates.

DnaD alanine substitution strains were generated by a blue/white screening assay using CW197 as parental strain and mutant plasmids obtained after Quickchange mutagenesis and sequencing as recombinant DNA (Fig. S3). X-gal 0.016% w/v was added to transformation plates for detection of β-galactosidase activity and selection of kanamycin resistant white colonies that integrated mutant DNA by double-recombination. Three individual white colonies per mutant were then streaked onto a medium either with or without IPTG to identify alleles of interest.

### Biological resources: E. coli strains and plasmids

*E. coli* transformations were performed in CW198 via heat-shock following the Hanahan method (48) for plasmids harbouring *dnaD* and propagated in LB with appropriate antibiotics at 37°C unless indicated otherwise in method details. Plasmids are listed in the Table S2 (sequences are available upon request). DH5α [F^-^Φ80*lac*ZΔM15 Δ(*lac*ZYA-*arg*F) U169 *rec*A1 *end*A1 *hsd*R17(r_k_^-^, m_k_^+^) *pho*A *sup*E44 *thi*-1 *gyr*A96 *rel*A1 λ^-^] (49) was used for other plasmids construction. HM1784 [BTH101 Δ*rnh*::*kan*] was used for bacterial-2-hybrid experiments and constructed by P1 transduction of Δ*rnh*::*kan* from JW0204 (Keio collection) into BTH101 [F-, *cya-99*, *ara*D139*, gal*E15*, gal*K16*, rps*L1 *(Str^r^)*, *hsd*R2*, mcr*A1*, mcr*B1 (Euromedex)]. Descriptions, where necessary, are provided below.

pCW123, pCW141, pCW142, pCW143, pCW153, pCW163, pCW214, pHM543, pHM544, pHM545 were generated by Quickchange mutagenesis using the oligonucleotides listed in Table S3.

pCW4 was generated by cloning *HindIII-SphI* PCR fragments generated using oligonucleotides listed in Table S3.

pDS84, pDS119, pDS120, pDS126, pDS127, pDS132, pHM359, pHM638, pHM640, pHM642, pHM644 were generated by cloning *Asp718I-BamHI* PCR fragments generated using the oligonucleotides listed in Table S3.

pCW66, pCW137, pCW171, pCW213, pSP075, pSP080, pSP081, pSP082, pSP083, pSP085 were generated by ligase-free cloning via two-step assembly processes using oligonucleotides listed in Table S3. The underlined part of each primer indicates the region used to form an overlap. FastCloning (50) was used with minor modifications. PCR products (15 μl from a 50 μl reaction) were mixed and then subjected to a heating/cooling regime: two cycles of 98°C for 2 minutes then 25°C for 2 minutes, then one cycle of 98°C for 2 minutes then 25°C for 60 minutes. After cooling *DpnI* restriction enzyme (1 μl) was added to digest parental plasmids and the mixtures were incubated at 37°C for ∼4 hours. Following digestion 10 μl of the PCR mixture was transformed into chemically competent *E. coli*. Where several primer pairs are listed for the construction of a single plasmid (Multi-step assembly column in Table S3), multiple rounds of ligase free cloning were performed to obtain the final constructs.

DnaD alanine-scan mutant plasmids were generated by Quickchange mutagenesis using oligonucleotides listed in Table S4. Cloning protocols were adapted to a 96-well plate format for PCR amplification of mutant plasmids, heat-shock and transformation recovery. All plasmids were sequenced.

### Biological resources: oligonucleotides

All oligonucleotides were purchased from Eurogentec. Oligonucleotides used for plasmid construction are listed in Table S3, those generated by the Quickchange program are listed in Table S4 and oligonucleotides used for qPCR are listed in Table S5. Details about the design of the Quickchange mutagenesis software can be found below.

Quickchange mutagenesis was used for the construction of the DnaD mutant plasmid library. Each point mutant was assembled by PCR using mutagenic primers carrying a single alanine substitution (51). We generated all mutant primer pairs via an in-house Quickchange program that optimised sequences according to key features in site-directed mutagenesis primer design (52). These include sequence length adjustments based on: (i) the melting temperature (T_M_) of the oligonucleotide part that anneals to the template plasmid, (ii) the T_M_ corresponding to a primer pair overlapping section, (iii) the GC-content within different sections of individual primers, (iv) the presence of a GC-clamp at every oligonucleotide 3’- end, and (v) the T_M_ difference between forward and reverse primer pairs. Code was written in Java and available online.

### Statistical analyses

Statistical analysis was performed using Student’s t-tests and p-values are given in figure legends. The exact value of *n* is given in method details and represents the number of biological repeats for an experiment. Tests were based on the mean of individual biological replicates and error bars indicate the standard error of the mean (SEM) across these measurements. Differences were considered as significant if their associated p-value was below 0.05. For sporulation experiments showing protein depletion from the origin, the standard error of the mean was propagated by addition of the error from individual terms.

### Data availability: new software

All original code to generate the QuickChange program has been deposited at Zenodo and is publicly available as of the date of publication. The DOI of the software is 10.5281/zenodo.5541537.

### Microscopy

To visualize cells by microscopy during the exponential growth phase, starter cultures were grown in imaging medium (Spizizen minimal medium supplemented with 0.001 mg/mL ferric ammonium citrate, 6 mM magnesium sulphate, 0.1 mM calcium chloride, 0.13 mM manganese sulphate, 0.1% w/v glutamate, 0.02 mg/mL tryptophan) with 0.5% v/v glycerol, 0.2% w/v casein hydrolysate and 0.1 mM IPTG at 37°C. Saturated cultures were diluted 1:100 into fresh imaging medium supplemented with 0.5% v/v glycerol and 0.1 mM IPTG and allowed to grow for three mass doublings. For DnaD mutants, early log cells were then spun down for 5 minutes at 9000 rpm, resuspended in the same medium lacking IPTG and further incubated for 90 minutes before imaging.

Cells were mounted on ∼1.4% w/v agar pads (in sterile ultrapure water) and a 0.13- to 0.17-mm glass coverslip (VWR) was placed on top. Microscopy was performed on an inverted epifluorescence microscope (Nikon Ti) fitted with a Plan Apochromat Objective (Nikon DM 100x/1.40 Oil Ph3). Light was transmitted from a CoolLED pE-300 lamp through a liquid light guide (Sutter Instruments), and images were collected using a Prime CMOS camera (Photometrics). The fluorescence filter sets were from Chroma: GFP (49002, EX470/40 (EM), DM495lpxr (BS), EM525/50 (EM)), mCherry (49008, EX560/40 (EM), DM585lprx (BS), EM630/75 (EM)) and DAPI (49000, EX350/50, DM400lp, EM460/50). Digital images were acquired using METAMORPH software (version 7.7) and analysed using Fiji software (53). All experiments were independently performed at least twice, and representative data are shown.

The number of origins was quantified using the Trackmate plugin within the Fiji software (54). Background was subtracted from fluorescence images set to detect 8-10 pixel blob diameter foci over an intensity threshold of 150 relative fluorescence units. A mask containing the detected origin foci was created and merged with the nucleoids channel, and the number of origins per nucleoid was determined and averaged for a minimum of 100 cells from each strain that was examined. This analysis was performed for at least two individual biological repeats.

### Phenotype analysis of dnaD mutants using the inducible dnaD-ssrA strain

Strains were grown for 18 hours at 37°C on NA plates unless otherwise stated (spot-titre assays) or in Penassay Broth (PAB, plate reader experiments) either with or without IPTG (0.1 mM). All experiments were independently performed at least twice and representative data are shown.

### Immunoblot analysis

Proteins were separated by electrophoresis using a NuPAGE 4-12% Bis-Tris gradient gel run in MES buffer (Life Technologies) and transferred to a Hybond-P PVDF membrane (GE Healthcare) using a semi-dry apparatus (Bio-rad Trans-Blot Turbo). DnaA, DnaD and FtsZ were probed with polyclonal primary antibodies (Eurogentec) and then detected with an anti-rabbit horseradish peroxidase-linked secondary antibody (A6154, Sigma) using an ImageQuant LAS 4000 mini digital imaging system (GE Healthcare). Detection of DnaA, DnaD and FtsZ was within a linear range. Experiments were independently performed at least twice and representative data are shown.

### Marker frequency analysis

Strains were grown in PAB overnight at 37°C and diluted 1:100 the next morning in PAB. Cells were allowed to grow for 4 hours at 37°C or incubated until they reached an optical density of 0.4 for cold-sensitive assays performed at 20°C. Five hundred microliter samples were harvested and immediately mixed with sodium azide (1% w/v final) to arrest growth and genome replication. Cultures were collected by centrifugation, the supernatant was discarded and pellets were flash frozen in liquid nitrogen before gDNA extraction via the DNeasy blood and tissue kit (Qiagen).

qPCR was performed using the Luna qPCR mix (NEB) to measure the relative amount of origin DNA compared to the terminus. PCR reactions were run in a Rotor-Gene Q Instrument (Qiagen) using serial dilutions of the DNA and spore DNA was used as a control. Oligonucleotide primers were designed to amplify *incC* (qSF19/qSF20) and the terminus (qPCR57/qPCR58), were typically 20–25 bases in length and amplified a ∼100 bp PCR product (Table S5). Individual *ori:Ter* ratios were obtained in three steps: first, every Ct value was normalised to ^1^/_2Ct_, the dilution factor used during the qPCR and technical triplicates were averaged to a single enrichment value; second, origin enrichment was normalised by corresponding terminus values; third, *ori:Ter* values were normalised by the enrichment obtained for spore DNA. Error bars indicate the standard error of the mean for 2-4 biological replicates.

### Protein structure representations

Protein representations were generated using the Pymol Molecular Graphics 2.1 software (55).

### Bacterial two-hybrid assays

*E. coli* strain HM1784 was transformed using a combination of complementary plasmids and grown to an OD_600nm_ of 0.5 in LB containing ampicillin and spectinomycin, before diluting 1:10,000 and spotting onto nutrient agar plates containing antibiotics and the indicator X-gal (0.008% w/v). Plates were incubated at 30°C for 24 to 48 hours and imaged using a digital camera. Experiments were independently performed at least twice and representative data are shown.

### Protein purification

Wild-type *dnaA* and *dnaD* were amplified by PCR using genomic DNA from *B. subtilis* 168CA and respectively cloned into pSF14 and pSF17 containing a His^14^-SUMO tag. DnaD mutants were created from pSF17 via quickchange reactions to introduce single or multiple substitutions. Plasmids were propagated in *E. coli* DH5α and transformed in BL21(DE3)- pLysS for expression. Strains were grown in LB medium at 37°C. Overnight cultures were diluted 1:100 the next morning and at A_600_ of 0.6, 1 mM IPTG was added before further incubation at 30°C for 4 hours. Cells were harvested by centrifugation at 7000 g for 20 minutes, DnaA expression pellets resuspended in 40 ml of DnaA Ni^2+^ Binding Buffer (30 mM HEPES [pH 7.6], 250 mM potassium glutamate, 10 mM magnesium acetate, 30 mM imidazole), DnaD pellets in 40 ml of DnaD Ni^2+^ Binding Buffer (40 mM Tris-HCl [pH 8.0], 0.5 M NaCl, 5% w/v glycerol, 1 mM EDTA, 20 mM imidazole), each containing 1 EDTA-free protease inhibitor tablet (Roche #37378900) and then flash frozen in liquid nitrogen. Cell pellet suspensions were thawed and incubated with 0.5 mg/ml lysozyme on ice for 1h before disruption by sonication (1 hour at 20 W with 20 seconds pulses/rests intervals). Cell debris were removed from the lysate by centrifugation at 24,000 g for 30 minutes at 4°C, then passed through a 0.2 µm filter for further clarification. Further purification steps were performed at 4°C using a FPLC with a flow rate of 1 ml/min.

Clarified lysates were applied to a 1 ml HisTrap HP column (Cytiva). For DnaA, an additional wash with 10 ml Ni^2+^ High Salt Wash Buffer (30 mM HEPES [pH 7.6], 1 M potassium glutamate, 10 mM magnesium acetate, 30 mM imidazole) was performed. Materials bound to the column were washed with 10 ml of 10% Ni^2+^ Elution Buffer (DnaA: 30 mM HEPES [pH 7.6], 250 mM potassium glutamate, 10 mM magnesium acetate, 1 M imidazole; DnaD: 40 mM Tris-HCl [pH 8.0], 0.5 M NaCl, 1 mM EDTA, 0.5 M imidazole) and proteins were eluted with a 10 ml linear gradient (10-100%) of Ni^2+^ Elution Buffer. For DnaA, fractions containing the protein were applied to a 1 ml HiTrap Heparin HP affinity column (Cytiva) equilibrated in H Binding Buffer (30 mM HEPES [pH 7.6], 100 mM potassium glutamate, 10 mM magnesium acetate) and elution was carried out with a 20 ml linear gradient (20–100%) of H Elution Buffer (30 mM HEPES [pH 7.6], 1 M potassium glutamate, 10 mM magnesium acetate). Fractions containing proteins of interest were pooled and digested with 10 µl of 10 mg/ml His^14^-Tev-SUMO protease (56). For DnaD, digestion was performed at room temperature over the course of 48 hours and the same amount of His^14^- Tev-SUMO protease was added after 24 hours digestion.

Digestion reactions were applied to a 1 ml HisTrap HP column to capture non-cleaved monomers, His^14^-SUMO tag and His^14^-TEV-SUMO protease. Cleaved proteins were collected in the flow-through and their purity was confirmed using SDS-PAGE. Glycerol was added (DnaA: 20% v/v final; DnaD: 10% v/v final) and proteins aliquots were flash frozen in liquid nitrogen before being stored at -80°C.

### SEC-MALS

Experiments were conducted on a system comprising a Wyatt HELEOS-II multi-angle light scattering detector and a Wyatt rEX refractive index detector linked to a Shimadzu HPLC system (SPD-20A UV detector, LC20-AD isocratic pump system, DGU-20A3 degasser and SIL-20A autosampler) and the assays performed at 20°C. Solvent was 0.2 µm filtered before use and a further 0.1 µm filter was present in the flow path. The column was equilibrated with at least 2 column volumes of 40 mM Tris-HCl [pH 8], 500 mM NaCl, 1 mM EDTA, 20 mM imidazole, 2.5% v/v glycerol before use and flow was continued at the working flow rate until baselines for UV, light scattering and refractive index detectors were all stable.

The sample injection volume was of 100 µl, the Shimadzu LabSolutions software was used to control the HPLC and the Astra 7 software for the HELEOS-II and rEX detectors. The Astra data collection was 1 minute shorter than the LC solutions run to maintain synchronisation. Blank buffer injections were used as appropriate to check for carry-over between sample runs. Data were analysed using the Astra 7 software. Molecular weights were estimated using the Zimm fit method with degree 1 and a value of 0.179 was used for protein refractive index increment (dn/dc).

### Oligomerisation crosslinking assay

Protein concentrations were adjusted to 300 nM in 1.2x Conjugation Buffer (20 mM HEPES [pH 7.2], 100 mM NaCl, 10 mM magnesium acetate) and 15 µl reactions were allowed to equilibrate at 20°C. BS_3_ (ThermoFisher A39266) was used as a crosslinking agent and diluted in water to a stock concentration of 750 µM. For final concentrations of 250 nM protein and 500x molar excess BS3, 3 µl of crosslinker were gently mixed with protein reactions. All reactions were incubated for 6 min at 20°C, then stopped by adding Quenching Buffer (final concentration 50 mM Tris-HCl [pH 7.5]) and further incubated for 10 min. NuPAGE loading dye (ThermoFisher NP0007) supplemented with excess DTT was added to all reactions and samples were fixed for 5 min at 95°C before loading on a 4-12% Bis-Tris gel (Invitrogen WG1402BOX). The same samples without BS_3_ were run as a loading control and stained using a coomassie protein stain (Abcam ab119211) because detection via our anti-DnaD antibody (Eurogentec) was not fully consistent between wild-type and variant alleles of *dnaD*. After migration, species were transferred to a Hybond-P PVDF membrane (GE Healthcare) using a semi-dry apparatus (Bio-rad Trans-Blot Turbo). Proteins were probed via α-DnaD polyclonal primary antibodies (Eurogentec) and then detected with an anti-rabbit horseradish peroxidase-linked secondary antibody (A6154, Sigma) using an ImageQuant LAS 4000 mini digital imaging system (GE Healthcare). Experiments were independently performed at least twice and representative data is shown.

### Sequence alignments

Multiple protein sequence alignments were performed using the Clustal Omega tool (57). Protein sequences are listed in Table S7.

### Phenotype analysis of dnaA mutants using the inducible oriN strain

Strains were grown for 48 hours at 30°C, 37°C or 48°C on NA plates either with or without IPTG (0.1 mM). All experiments were independently performed at least twice and representative data is shown.

### DNA strand separation assay

DNA scaffolds that contained one oligonucleotide labelled with BHQ2, one with Cy5 and one unlabelled (12.5 nM final concentration) were diluted in 10 mM HEPES-KOH (pH 8), 100 mM potassium glutamate, 2 mM magnesium acetate, 30% glycerol, 10% DMSO and 1 mM nucleotide (ADP or ATP). All the reactions were prepared on ice to ensure the stability of the DNA probe, then allowed to equilibrate at 20°C. DnaA was added to a final concentration of 650 nM to allow displacement of all the probes. Reactions were performed using a flat-bottom black polystyrene 96-well plate (Costar #CLS3694) in triplicate and fluorescence was detected every minute over 60 min with a plate reader (BMG Clariostar). For all reactions a negative control without protein was used as background. At each timepoint the average background value was subtracted from the experimental value, thus reporting the specific DnaA activity on a single substrate. Error bars indicate the standard error of the mean over three biological replicates.

### ChIP-qPCR

Chromatin immunoprecipitation and quantitative PCR were performed as previously described (58) with minor modifications detailed below.

Strains were grown overnight at 30°C in Spizizen salts supplemented with tryptophan (20 µg/ml), glutamate (0.1% w/v), glucose (0.5% w/v) and casamino acid (0.2% w/v). The following day cultures were diluted 1:100 into fresh medium and allowed to grow to an A_600_ of 0.4. Samples were resuspended in PBS and cross-linked with formaldehyde (final concentration 1% v/v) for 10 minutes at room temperature, then quenched with 0.1 M glycine. Cells were pelleted at 4°C, washed three times with ice-cold PBS (pH 7.3) then frozen in liquid nitrogen and stored at -80°C. Frozen cell pellets were resuspended in 500 µl of lysis buffer (50 mM NaCl, 10 mM Tris-HCl pH 8.0, 20% w/v sucrose, 10 mM EDTA, 100 µg/ml RNase A, ¼ complete mini protease inhibitor tablet (Roche), 2000 K U/µl Ready-Lyse lysozyme (Epicentre)) and incubated at 37°C for 30 min to degrade the cell wall. 500 µl of immunoprecipitation buffer (300 mM NaCl, 100 mM Tris-HCl pH 7.0, 2% v/v Triton X-100, ¼ complete mini protease inhibitor tablet (Roche), 1 mM EDTA) was added to lyse the cells and the mixture was incubated at 37°C for a further 10 minutes before cooling on ice for 5 minutes. DNA samples were sonicated (40 amp) three times for 2 minutes at 4°C to obtain an average fragment size of ∼500 to 1000 base pairs. Cell debris were removed by centrifugation at 4°C and the supernatant transferred to a fresh Eppendorf tube. To determine the relative amount of DNA immunoprecipitated compared to the total amount of DNA, 100 µl of supernatant was removed, treated with Pronase (0.5 mg/ml) for 60 minutes at 37°C then stored on ice. To immunoprecipate protein-DNA complexes, 800 µl of the remaining supernatant was incubated with rabbit polyclonal anti-DnaA, anti-DnaD and anti- DnaB antibodies (Eurogentec) for 1 hour at room temperature. Protein-G Dynabeads (750 µg, Invitrogen) were equilibrated by washing with bead buffer (100 mM Na_3_PO_4_, 0.01% v/v Tween 20), resuspended in 50 µl of bead buffer, and then incubated with the sample supernatant for 1 hour at room temperature. The immunoprecipated complexes were collected by applying the mixture to a magnet and washed with the following buffers for 15 minutes in the respective order: once in 0.5X immunoprecipitation buffer; twice in 0.5X immunoprecipitation buffer + NaCl (500 mM); once in stringent wash buffer (250 mM LiCl, 10 mM Tris-HCl pH 8.0, 0.5% v/v Tergitol-type NP-40, 0.5% w/v sodium deoxycholate 10 mM EDTA). Finally, protein-DNA complexes were washed a further three times with TET buffer (10 mM Tris-HCl pH 8.0, 1 mM EDTA, 0.01% v/v Tween 20) and resuspended in 100 µl of TE buffer (10 mM Tris-HCl pH 8.0, 1 mM EDTA). Formaldehyde crosslinks of both the immunoprecipate and total DNA were reversed by incubation at 65°C for 16 hours in the presence of 1,000 U Proteinase K (excess). The reversed DNA was then removed from the magnetic beads, cleaned using QIAquick PCR Purification columns (Qiagen) and used for qPCR analysis.

qPCR was performed using the Luna qPCR mix (NEB) to measure the amount of genomic loci bound to DnaA, DnaD and DnaB. PCR reactions were run in a Rotor-Gene Q Instrument (Qiagen) using serial dilutions of the immunoprecipitate and total DNA control as template. Oligonucleotide primers were designed to amplify *oriC* (qSF11/qSF12), *oriN* (qSF5/qSF6) and the non-specific locus *yhaX* (oWKS145/oWKS146 (39)), and were typically 20–25 bases in length and amplified a ∼100 bp PCR product (Table S5). Error bars indicate the standard error of the mean for 6-8 biological replicates.

### Pull-down assay of His_6_-DnaA^DI^-DnaD^NTD^ complexes

BL21 (DE3) *E. coli* cells containing the different expression plasmids (pSP075, pSP080, pSP081, pSP082, pSP083 and pSP085) were grown overnight in 5 ml of LB supplemented with kanamycin at 37°C. The following day cells were diluted in 50 ml of fresh medium until A_600_ reached 0.5. Protein expression was induced by adding 1 mM IPTG for 4 hours at 30°C. Cells were collected by centrifugation and resuspended in 2 ml of resuspension buffer (30 mM Hepes pH 7.5, 250 mM potassium glutamate, 10 mM magnesium acetate, 20% w/v sucrose, 30 mM imidazole) supplemented with 1 EDTA-free protease inhibitor tablet. Bacteria were lysed with two sonication cycles at 10 W for 3 minutes with 2 second pulses. Cell debris were pelleted by centrifugation at 25,000 g at 4°C for 30 minutes and the supernatant was filtered through 0.2 µm filters. The clarified lysate was then loaded onto Ni-NTA spin columns (QIAgen) and proteins purified according to manufacturer protocol washing the column with Washing Buffer (30 mM Hepes pH 7.5, 250 mM potassium glutamate, 10 mM magnesium acetate, 20% sucrose, 100 mM imidazole) and eluting bound proteins with elution buffer (30 mM Hepes pH 7.5, 250 mM potassium glutamate, 10 mM magnesium acetate, 20% w/v sucrose, 1 M imidazole). The eluates were loaded on a NuPAGE 4-12% Bis-Tris gradient gel run in MES buffer (Life Technologies) and analysed with IstantBlue staining (Merck).

## RESULTS

DnaD is an essential DNA replication initiation protein in the model organism *B. subtilis* and in opportunistic human pathogens such as *Staphylococcus aureus* and *Streptococcus pneumoniae* (59–61). However, the specific activities required by DnaD at the chromosome origin *in vivo* are unclear. To address these questions, we sought to identify essential amino acids in *B. subtilis* DnaD and then to determine the function of each essential residue.

### Identification of essential residues in DnaD

Functional analysis of bacterial DNA replication initiation proteins *in vivo* is challenging because they are required for viability. Mutation of an essential feature will be lethal, while mutations that severely disable function can result in the rapid accumulation of compensatory suppressors. To circumvent these issues, a bespoke inducible complementation system was developed for *dnaD* (*P_HAT_*-*dnaD-ssrA*, Fig. S2). Upon repression of the ectopic *dnaD-ssrA*, the functionality of *dnaD* alleles at the endogenous locus can be determined.

A plasmid for allelic exchange of the endogenous *dnaD* gene was created (Fig. S3). Using this template, a library of 222 single alanine substitution mutants (all codons save for the start, stop and naturally occurring alanine) was generated and sequenced. To ensure mutagenesis of the native *dnaD* following transformation, a recipient strain was constructed containing both the *dnaD* operon replaced by *bgaB* (encoding the enzyme β-galactosidase) and the inducible *dnaD-ssrA* complementation system (Fig. S3). Thus, replacement of *bgaB* with *dnaD* mutants can be detected on selective media supplemented with a chromogenic substrate (e.g. white colonies, Fig. 1B) and confirmed by chloramphenicol sensitivity.

Following construction of the *dnaD* alanine substitution library, cultures grown in a microtiter plate were monitored in a plate reader to assess the functionality of each mutant. The data revealed growth defects for several alanine substitutions, spread throughout the protein (Fig. 1C-F). Immunobloting was used to determine the expression of thirteen DnaD variants conferring a lethal phenotype, and this analysis showed that five were not detected (Fig. S4). Therefore, for the least ambiguous interpretation of results, we focussed on essential alanine substitutions that were expressed near the wild-type level. Taken together, this analysis identified fourteen alanine substitutions in DnaD that retained detectable protein expression and produced a growth phenotype, nine of which were essential for viability (Figs. 1D-F and S4, Table S6).

### DnaD essential mutants abolish DNA replication initiation

While essential for DNA replication initiation in *B. subtilis*, DnaD has also been implicated in other key cellular processes including replication restart, chromosome organization, and DNA repair (39,62–65). To ascertain whether *dnaD* alanine mutants were specifically impaired in DNA replication initiation, we characterized chromosome content in these strains using fluorescence microscopy.

During steady-state growth in defined minimal media, wild-type *B. subtilis* cells typically display a pair of chromosome origins per nucleoid (*ori:nuc*), each orientated towards one cell pole (66). In contrast, when chromosome replication is inhibited, nucleoids typically contain a single *oriC* signal located near the centre of the bulk DNA (Fig. 2A) (67). Therefore, to evaluate the impact of *dnaD* mutants on DNA replication, a strain was constructed with a fluorescently labelled nucleoid associated protein (Hbs-GFP) to detect the chromosome (68) and a fluorescent reporter-operator system (*tetO* array with TetR-mCherry) (69) to detect the relative number of *oriC* regions in every cell (Fig. 2B). Cells were imaged following repression of the ectopic *dnaD-ssrA* for 90 minutes. All *dnaD* alanine mutants produced a phenotype characteristic of non-replicating chromosomes, with well separated nucleoids often containing a single central TetR-mCherry focus (Fig. 2C). Quantitative analysis of microscopy images showed that all mutants appeared to display fewer origins and a decrease in the number of nucleoids per cell (Fig. 2D), resulting in a reduction of the *ori:nuc* ratio compared to wild-type DnaD (Fig. 2E). These results indicate that the essential DnaD amino acids are required to initiate DNA replication *in vivo*.

**Figure 2.**
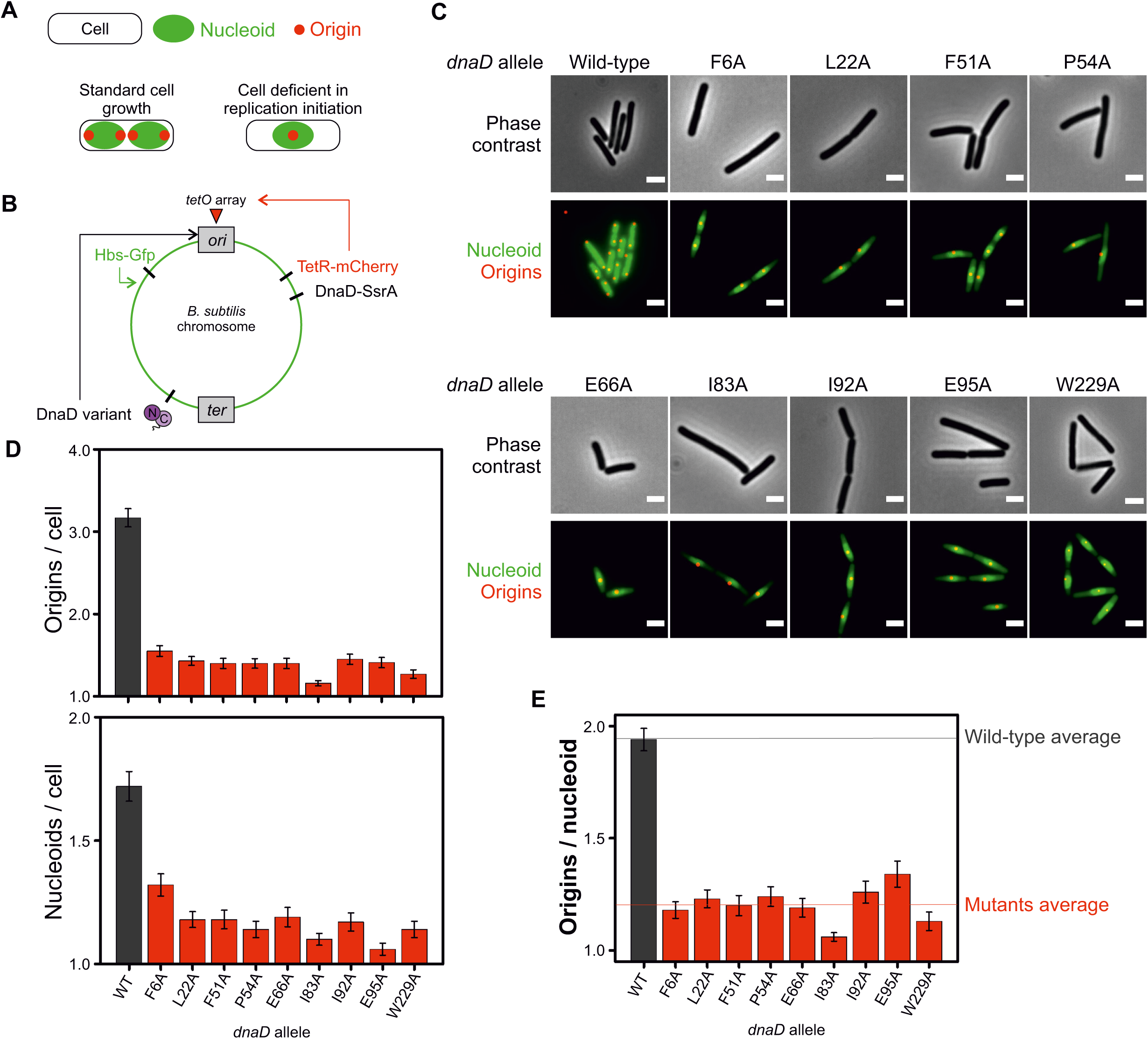
Essential DnaD mutants are defective in DNA replication initiation. **(A)** Schematics of the number of origins and nucleoids per cell showing the typical patterns associated with cells displaying a normal growth phenotype or cells that are deficient in replication initiation. **(B)** Schematics of the dual fluorescence system engineered to label chromosome origins and the nucleoid. TetR-mCherry (red fluorescence) is constitutively expressed and binds an array of *tetO* sites located near the origin of replication. Hbs-GFP (green fluorescence) is constitutively expressed, non-specifically binds double-stranded DNA and allows visualisation of the nucleoid. **(C)** Representative images of *dnaD* essential substitutions observed by fluorescence microscopy via the system described in (A). Red dots show chromosome origins and the green signal allows localisation of the nucleoid. Wild-type corresponds to a strain encoding the ectopic dnaD-ssrA cassette that was depleted from IPTG and relied on the endogenous copy of wild-type *dnaD* (CW517). F6A (CW540), L22A (CW536), F51A (CW534), P54A (CW539), E66A (CW538), I83A (CW533), I92A (CW537), E95A (CW532), W229A (CW535). **(D)** Number of origins per cell (top) and nucleoids per cell (bottom) corresponding to the single-cell image analysis on DnaD variants from the experiments shown in (C). **(E)** Origin per nucleoid ratios corresponding to the single-cell image analysis performed on the data obtained from experiments shown in (C). Error bars in (D-E) show the standard error of the mean for at least two biological replicates where over 100 cells were counted for *dnaD* wild- type and mutant backgrounds.

### A DnaD tetramer is necessary for DNA replication initiation in vivo

The crystal structure of DnaD^NTD^ has been solved as a symmetric homodimer, while biochemical experiments and structural modelling suggest assembly into a tetramer or higher-order oligomer (41, 42). Critically, the active form of DnaD *in vivo* was not known. The alanine scan showed that replacement of either Phe6 or Leu22 was lethal (Figs. 1D, 3A) and immunobloting indicated that these DnaD variants are stable *in vivo* (Fig. 3B). Mapping the residues onto the DnaD^NTD^ crystal structure revealed that Leu22 is buried within the proposed dimerization interface while Phe6 is exposed towards the predicted dimer:dimer interface (Fig. 3C)(41). Note that Figure 3C also highlights the positions of alanine substitution variants that, while viable, were observed to altered colony morphology and to decrease the frequency of DNA replication initiation (Fig. S5). The location of these residues suggests they may play a role in DnaD homo-oligomerisation.

**Figure 3.**
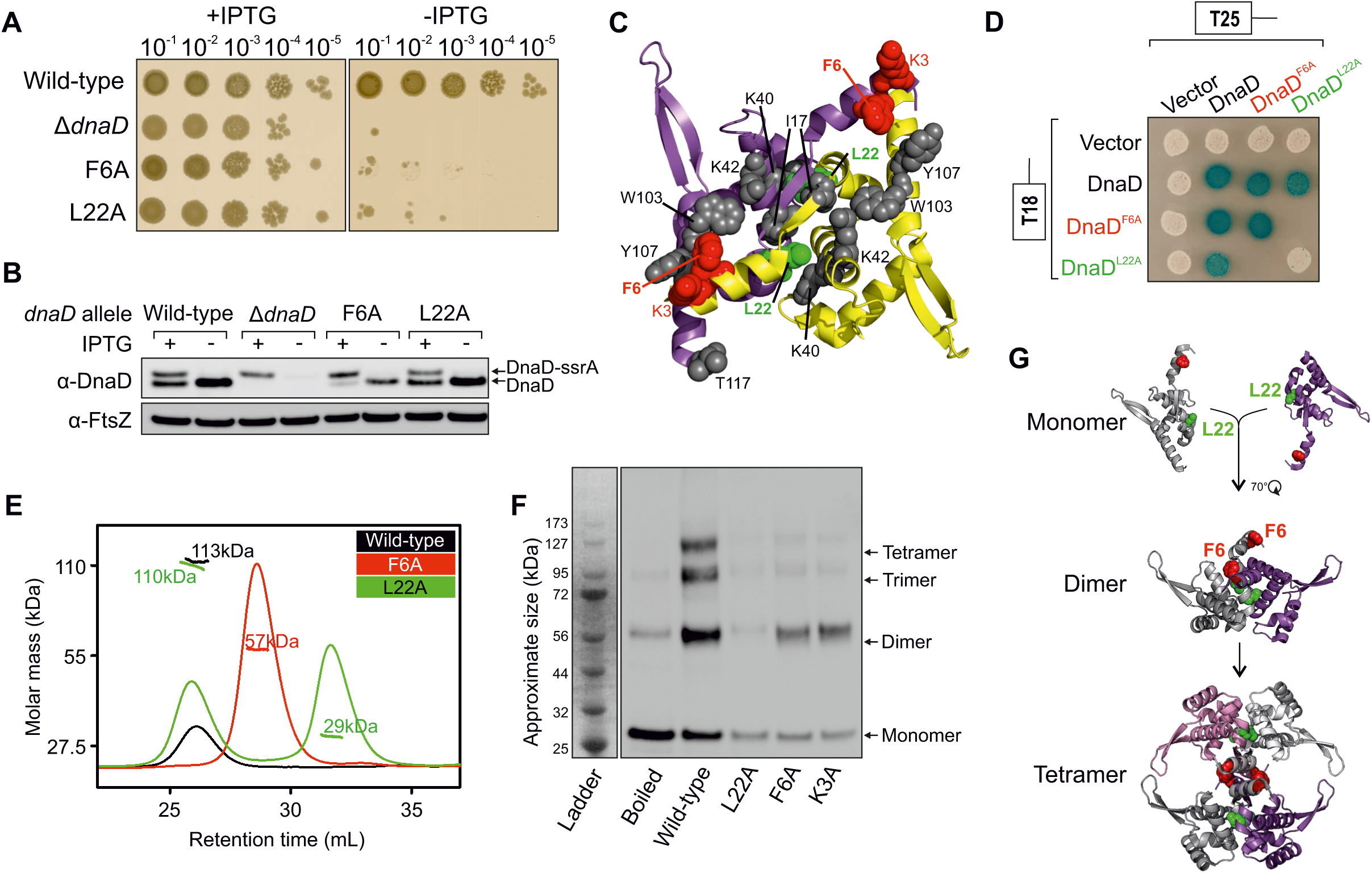
Lethal alanine substitutions in DnaD disrupt tetramer formation. **(A)** DnaD N-terminal domain crystal structure (PDB 2v79) showing an extensive interface mapped with residues that produced a different phenotype than wild-type after the alanine scan. Purple and yellow subunits indicate individual protomers. Residues are marked as red for lethal and grey for altered growth (translucent/heterogeneous phenotype or slow growth). **(B)** Spot titre analysis showing that substitutions *dnaD^F6A^* (CW310) and *dnaD^L22A^* (CW292) produced a lethal phenotype *in vivo.* The presence or absence of IPTG indicates the induction state of the ectopic *dnaD-ssrA* cassette. Wild-type (CW162), Δ*dnaD* (CW197). **(C)** Immunoblot showing that DnaD^F6A^ and DnaD^L22A^ are expressed *in vivo* and remain stable upon depletion of the ectopic DnaD-SsrA copy (no IPTG condition). Detection of the tubulin homolog FtsZ was used as a loading control. Strains are the same as used in (B). **(D)** Bacterial two-hybrid assay showing the effect of mutants DnaD^L22A^ and DnaD^F6A^ on self-interaction. White spots indicate a lack of interaction and blue spots the interaction between two protein variants. White colonies observed with empty vectors indicate that the detected interactions are specific. Plasmids used in this assay are listed in Table S2. **(E)** SEC-MALS analysis of purified DnaD protein variants. The UV spectrum (continuous lines) was normalised as a relative refraction index and the molar mass corresponding to each protein is shown as shorter/thicker lines overlapping the different peaks. Masses corresponding to each peak are annotated on the plot. **(F)** Immunoblot following migration and transfer of BS^3^ crosslinked DnaD species using SDS-PAGE. The ladder indicates the approximate size of fragments. The “Boiled” lane corresponds to a wild-type DnaD sample that was boiled prior to crosslinking. **(G)** Schematics of DnaD N-terminal domain oligomerisation pathway involving key residues F6 and L22. DnaD^L22^ allows formation of a dimer and DnaD^F6^ contributes to further assembly into a tetramer.

To begin assessing the DnaD self-interaction, a bacterial two-hybrid assay was employed. Full-length *dnaD* alleles were fused to catalytically complementary fragments of the *Bordetella pertussis* adenylate cyclase (T25 and T18) (70). Two-hybrid analysis showed that wild-type DnaD and DnaD^F6A^ self-interact, whereas DnaD^L22A^ has lost this capability (Fig. 3D). All DnaD proteins reported a positive interaction with wild-type DnaD, indicating that they were being stably expressed in the heterologous host (Fig. 3D). The lack of self-interaction observed for the DnaD^L22A^ variant suggests that Leu22 is key residue involved in DnaD dimerisation, consistent with previous structural data (41).

To further interrogate the quaternary structure of DnaD, we purified DnaD^L22A^ and DnaD^F6A^ and characterised these variants by size exclusion chromatography (SEC) (71) followed by multiple angle light scattering (MALS) (72). Wild-type DnaD was observed to run as a tetramer of approximately 113 kDa (theoretical molecular weight of 110 kDa) (Figs. 3E and S6A). SEC-MALS analysis showed that >50% of DnaD^L22A^ dissociated into a 29 kDa monomer, whereas DnaD^F6A^ was eluted exclusively as 57 kDa species, consistent with the protein forming a homodimer (Figs. 3E and S6A). It is unsurprising that DnaD^L22A^ retains some ability to assemble into a tetramer (Fig. 3E), as the substitution of leucine for alanine would not be expected to sterically disrupt the extensive dimer interface (Fig. 3C) (41). Consistent with SEC-MALS, the dimerization capability DnaD^L22A^ was found to be poor using the amine-specific crosslinker bis(sulfosuccinimidyl)suberate (BS^3^), while DnaD^F6A^ was competent to form a dimer but not to form a tetramer (Fig. 3F). Returning to the *dnaD* alanine scan, we appreciated that *dnaD^K3A^*was also lethal, albeit poorly expressed *in vivo* (Fig. S4). Nonetheless, recombinant DnaD^K3A^ could be purified, and BS^3^ crosslinking showed that this variant was also competent to form a dimer but not a tetramer (Figs. 3F and S6F).

Taken all together, the data indicate that DnaD oligomerization is mediated by the N-terminal domain, with Leu22 required for dimerisation and Lys3/Phe6 required for tetramerization (Fig. 3G). Moreover, the results suggest that adopting a tetrameric quaternary state is necessary to support DNA replication initiation *in vivo*.

### The interaction between DnaD^CTT^ and DnaB is necessary for DNA replication initiation in vivo

The alanine scan indicated that a cluster of residues in the unstructured C-terminal tail of DnaD are critical for cell growth, particularly Trp229 which is essential and stably expressed (Figs. 4A-C). Phylogenetic analysis indicates that Trp229 is conserved in species harbouring both *dnaD* and *dnaB*, but not *dnaD* alone (Fig. 4D and Table S7), suggesting that the DnaD^CTT^ could be an interaction site for DnaB.

**Figure 4.**
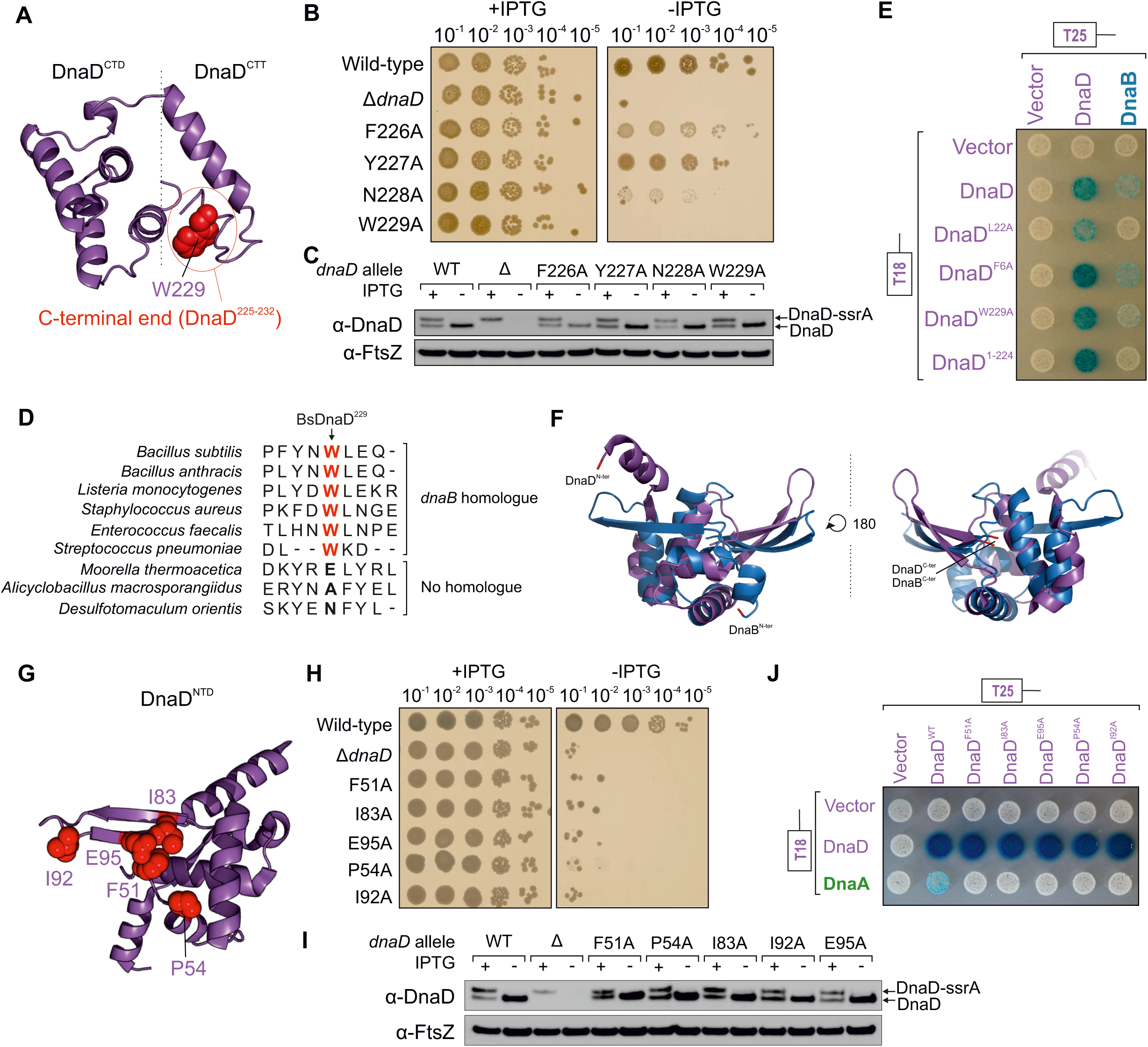
Lethal alanine substitutions in DnaD disrupt interactions with DnaB and DnaA. **(A)** Essential residues of DnaD, required for the interaction with DnaB, mapped onto a DnaD^CTD/CTT^ model. **(B)** Spot titre analysis showing that substitutions in DnaD^CTT^ variably affect cell growth *in vivo.* The presence or absence of IPTG indicates the induction state of the ectopic *dnaD-ssrA* cassette. Wild-type (CW162), Δ*dnaD* (CW197), F226A (CW284), Y227A (CW317), N228A (CW285), W229A (CW286). **(C)** Immunoblot showing that DnaD variants shown in (B) are expressed *in vivo* and remain stable upon depletion of the ectopic DnaD-SsrA copy (no IPTG condition). Detection of the tubulin homolog FtsZ was used as a loading control. Strains are the same as used in (B). **(D)** Protein sequence alignment of DnaD homologs showing that *B. subtilis* DnaD^W229^ is particularly conserved in species that also harbour a copy of DnaB. **(E)** Bacterial two-hybrid assay showing loss of interaction between DnaD variants and DnaB in the context of full-length proteins. White spots indicate a lack of interaction and blue spots the interaction between two protein variants. White colonies observed with empty vectors indicate that the detected interactions are specific. Plasmids used in this assay are listed in Table S2. **(F)** Structural alignment showing that the N-terminal domains of DnaD (PDB 2v79) and DnaB (PDB 5wtn) significantly overlap. **(G)** Essential residues of DnaD, required for the interaction with DnaA, mapped onto the DnaD^NTD^ crystal structure (PDB 2v79). **(H)** Spot titre analysis showing that substitutions in DnaD^NTD^ shown in (G) produced a lethal phenotype *in vivo.* The presence or absence of IPTG indicates the induction state of the ectopic *dnaD-ssrA* cassette. Wild-type (CW162), Δ*dnaD* (CW197), F51A (CW174), P54A (CW295), I83A (CW170), I92A (CW297), E95A (CW166). **(I)** Immunoblot showing that DnaD variants shown in (H) are expressed *in vivo* and remain stable upon depletion of the ectopic DnaD-SsrA copy (no IPTG condition). Detection of the tubulin homolog FtsZ was used as a loading control. Strains are the same as used in (H). **(J)** Bacterial two-hybrid assay showing loss of interaction between DnaD^NTD^ variants and DnaA in the context of full-length proteins. White spots indicate a lack of interaction and blue spots the interaction between two protein variants. White colonies observed with empty vectors indicate that the detected interactions are specific. Plasmids used in this assay are listed in Table S2.

To test this hypothesis, two-hybrid analysis was used to probe for protein:protein interactions. However, it was previously reported that interactions between full-length *B. subtilis* DNA replication initiation proteins could not be detected (31), consistent with the observation that expression of *B. subtilis* initiator proteins in *E. coli* is toxic (73, 74). We hypothesised that this phenotype arises from *B. subtilis* proteins interfering with DNA replication initiation in *E. coli* (e.g. *B. subtilis* DnaA binds to *E. coli* DnaA-boxes but cannot unwind *E. coli oriC*) (74). To circumvent these challenges and allow bacterial two-hybrid analysis of full-length *B. subtilis* DNA replication initiation proteins, we constructed a derivative of the *E. coli* two-hybrid strain with a deletion of the *rnhA* gene (encoding for RNase HI). RNase HI degrades stable DNA-RNA hybrid structures such as R-loops (75). The knockout of *rnhA* in *E. coli* allows formation of R-loops, which can be used to initiate DNA replication independently of DnaA and *oriC* (76).

Using this system, an interaction between wild-type DnaD and DnaB proteins was detected (Fig. 4E). In contrast, either altering Trp229 to alanine or deleting the last eight amino acids of DnaD reduced and disrupted the interaction with DnaB, respectively (Fig. 4E). It was also observed that the monomeric DnaD^L22A^ variant was unable to interact with DnaB, whereas the dimeric DnaD^F6A^ retained this capability (Fig. 4E). All DnaD variants retained a self-interaction with the wild-type protein, showing that they were being stably expressed (Fig. 4E). These results indicate that the interface between the distal end of DnaD^CTT^ and DnaB is essential for DNA replication initiation *in vivo*, and they suggest that DnaB recognition requires DnaD assembly into a homodimer.

Previous studies using protein truncation variants indicated that the DnaD^NTD^ interacts with DnaB^NTD^ (31). The observation that mutations in the DnaD^CTT^ abolish the interaction with DnaB, while leaving the DnaD^NTD^ intact, suggests that different DnaD:DnaB complexes are being detected in these assays. We also note that the N-terminal domains of DnaD and DnaB share structural homology (Fig. 4F) and that both promote dimerization/tetramerization (36, 37), such that the truncated variants used in previous studies (31) may be able to interact differentially in a two-hybrid assay.

### The interaction between DnaD^NTD^ and DnaA is necessary for DNA replication initiation in vivo

Models for the interaction between DnaD and DnaA have been proposed based on binding experiments using truncated proteins. These studies indicated that residues in the DnaD^NTD^ (31) and the DnaD^CTD^ (34) each contributed to DnaA binding. From the alanine scan it was observed that three of the residues at the proposed DnaA interface of the DnaD^NTD^ are essential (Phe51, Ile83, Glu95). Additionally, we identified two other lethal and stably expressed substitutions (DnaD^P54A^ and DnaD^I92A^) that mapped near these sites on the structure (Fig. 4G-I), suggesting they could also be involved in forming the DnaA interface.

In contrast, none of the residues in the proposed DnaA binding region of the DnaD^CTD^ were found to be essential (Fig. S7A-C). To investigate whether the interface between DnaD^CTD^ and DnaA was robust and single alanine substitutions were insufficient to disrupt binding, DnaD variants encoding multiple alanine substitutions were constructed (dnaD*^L129A/I132A^*, dnaD*^Y130A/E134A/E135A^*, dnaD*^I132A/E134A/E135A^*, dnaD*^K164A/K168A/E169A/V171A^*) (Fig. S7D-G). However, all of these *dnaD* alleles were viable and supported normal growth (Fig. S7C). These results indicate that the previously identified interface between DnaA and the DnaD^CTD^ (34) is not essential for DNA replication initiation *in vivo*. The DnaD^CTD^-DnaA interaction could play an auxiliary role that assists DnaD binding DnaA, or it could become important during certain environmental conditions or cell stresses via an alternative pathway. Two-hybrid analysis was used to investigate whether lethal alanine substitutions in DnaD^NTD^ perturb the interaction with DnaA. While an interaction between the full-length wild-type DnaD and DnaA proteins was detected, alanine substitutions within the proposed DnaD^NTD^ interface for DnaA abrogated the two-hybrid signal (Fig. 4J). All DnaD^NTD^ variants retained the ability to self-interact, indicating that they were being stably expressed (Fig. 4J). These results confirm and extend previous reports showing an interaction between DnaA and DnaD^NTD^ (31), and they show for the first time that this interaction is essential for DNA replication initiation *in vivo*.

### The interaction between DnaA^DI^ and DnaD is necessary for DNA replication initiation in vivo

Having identified the essential surface of DnaD required to interact with DnaA, we next determined essential residues in DnaA required to interact with DnaD. Previous two-hybrid and NMR studies implicated residues on the surface of DnaA^DI^ that interact with DnaD (31, 34). To investigate the physiological relevance of the proposed DnaA^DI^ interface, we replaced the endogenous *dnaA* gene with mutants encoding alanine substitutions at key residues (*dnaA^T26A^*, *dnaA^W27A^*, *dnaA^F49A^*) (Fig. 5A).

**Figure 5.**
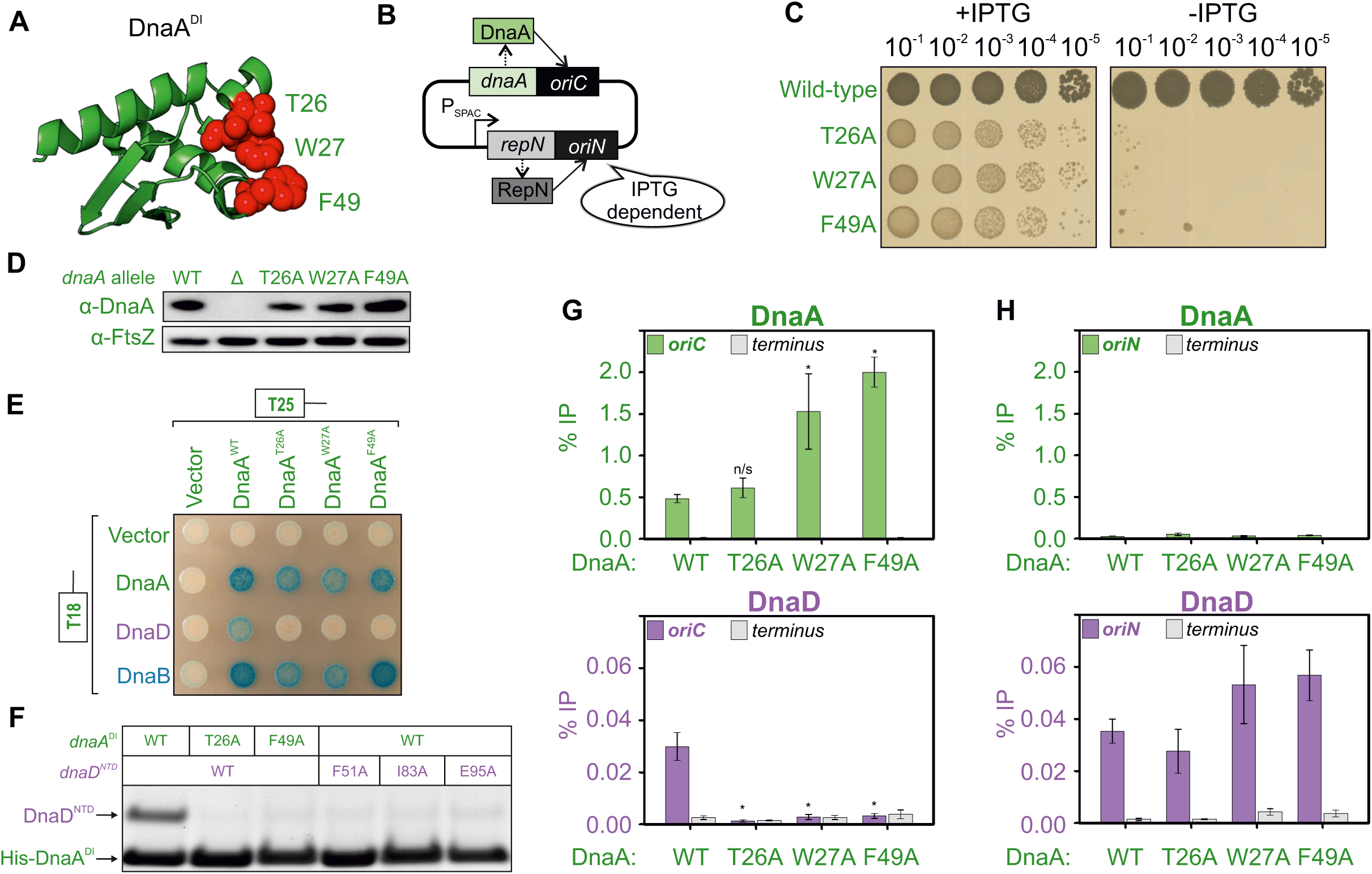
The interaction between DnaA domain I and DnaD^NTD^ is essential for DnaD recruitment to *oriC*. **(A)** Essential residues of DnaA, required for the interaction with DnaD, mapped onto the DnaA^DI^ crystal structure (PDB 4tps). **(B)** Schematics of the inducible *repN/oriN* system used to bypass mutations affecting DnaA activity in *B. subtilis*. Replication via *oriN* is turned on and off in the presence and absence of IPTG, respectively. **(C)** Spot titre analysis showing that DnaA^DI^ variants produced a lethal phenotype *in vivo*. The presence or absence of IPTG indicates the induction state of the *repN/oriN* system. Wild-type (HM1108), T26A (HM1540), W27A (HM1541), F49A (HM1542). **(D)** Immunobloting shows that DnaA^DI^ variants were expressed at a similar level to wild-type in the context of the *oriN* strain. Detection of the tubulin homolog FtsZ was used as a loading control. Wild-type (HM1524), Δ (HM1424), T26A (HM1537), W27A (HM1538), F49A (HM1539). **(E)** Bacterial two-hybrid assay showing the loss of interaction between DnaA variants and DnaD in the context of full-length proteins. The interaction between DnaA and DnaB was unaffected by DnaA^DI^ variants. White spots indicate a lack of interaction and blue spots the interaction between two protein variants. White colonies observed with empty vectors indicate that the detected interactions are specific. **(F)** Pull-down assay (stained SDS-PAGE) showing loss of interaction between His -DnaA^DI^ and DnaD^NTD^ when using variants of DnaA or DnaD. Plasmids used in (E-F) are listed in Table S2. **(G)** ChIP analysis showing that DnaA protein variants (wild-type, T26A, W27A and F49A, respectively HM949, HM1537, HM1538 and HM1539) remain enriched at *oriC*, whereas DnaD is not recruited to *oriC* when using DnaA^DI^ mutants. Primers used for the origin anneal within the *incC* region. * shows a p-value < 0.02 and n/s a non- significant difference. **(H)** Control for the ChIP analysis shown in (G). DnaA protein variants (wild-type, T26A, W27A and F49A) are not recruited to *oriN* while DnaD remains enriched at this site. Samples are the same as used in (G) with primers annealing within *oriN*.

To enable identification of essential amino acid residues without selecting for suppressor mutations, we utilized a strain in which DNA replication can initiate from a plasmid origin *(oriN*) integrated into the chromosome (Fig. 5B) (19). The activity of *oriN* requires its cognate initiator protein RepN, and both factors act independently of *oriC*/DnaA (note the RepN/*oriN* system also requires DnaD and DnaB for function) (Fig. S8) (77). Expression of *repN* was placed under the control of an IPTG-inducible promoter, thus permitting both the introduction of mutations into *dnaA* and their subsequent analysis following removal of the inducer to repress *oriN* activity. Cultures were grown overnight in the presence of IPTG and then serially diluted onto solid media with and without the inducer. The results show that the *dnaA^T26A^*, *dnaA^W27A^* and *dnaA^F49A^*mutants all inhibited colony formation, independent of temperature (Figs. 5C and S9A-B). Immunoblot analysis confirmed that all of the DnaA variants were expressed at a level similar to wild-type (Fig. 5D). This indicates that residues Thr26, Trp27 and Phe49 are essential for DnaA activity *in vivo*.

Two-hybrid analysis confirmed that alanine substitutions in DnaA at either Thr26, Trp27 or Phe49 inhibit the interaction with DnaD (Fig. 5E). These DnaA^DI^ variants retained the ability to interact with wild-type DnaA protein and with DnaB, indicating that they were stably expressed (Fig. 5E). The data suggest that the essential residues Thr26, Trp27, Phe49 of DnaA are required for the interaction with DnaD (Fig. 5E).

To investigate whether DnaA^DI^ and DnaD^NTD^ were sufficient to form a complex, pull-down assays between protein domains His_6_-DnaA^DI^ and DnaD^NTD^ were performed (Fig. S9C). Following co-expression of His_6_-DnaA^DI^ and DnaD^NTD^ in *E. coli*, cells were lysed and His_6_-DnaA^DI^ was captured using an immobilized nickel affinity chromatography spin column. While the wild-type DnaD^NTD^ was able to bind wild-type His_6_-DnaA^DI^, alanine substitutions in either protein greatly reduced the retention of DnaD^NTD^ (Fig. 5F). Analysis of cell lysates by SDS-PAGE revealed that all protein domains were being overexpressed to similar levels (Fig. S9D) and immunoblotting confirmed the identity of each polypeptide (Fig. S9E). These studies support and extend the previously proposed model for DnaA^DI^ interacting with DnaD^NTD^ (31), and they show for the first time that this interaction is essential for DNA replication initiation *in vivo*.

*The interaction of DnaA^DI^ with DnaD^NTD^ is required to recruit DnaD to the chromosome origin* DnaA binding to *oriC* is required to detect the enrichment of DnaD at the replication origin (32), therefore we wondered whether the DnaA^DI^-DnaD^NTD^ interaction described above was essential for DnaD recruitment. To test this hypothesis, we employed chromatin immunoprecipitation followed by quantitative PCR (ChIP-qPCR). Here, to support growth of lethal *dnaA* mutants, a strain harbouring a constitutively active version of *oriN* within the chromosome was used (78). In all cases, DnaA^DI^ variants remained specifically enriched at *oriC* while recruitment of DnaD was abolished (Fig. 5G). Consistent with previous studies (32, 79), the DnaA variants were observed to accumulate at *oriC* when DNA replication was perturbed. Note that in these strains DnaD remained enriched at *oriN,* as expected (Fig. 5H) (39).

The ChIP data are consistent with the model that an essential function of the DnaA^DI^- DnaD^NTD^ interaction is to recruit DnaD to *oriC*. However, an essential activity of DnaA is also to unwind *oriC*, and it has been suggested that DnaA^DI^ has an affinity for ssDNA {Abe, 2007 #2368}. It was conceivable that a defect in *oriC* unwinding might indirectly impact DnaD recruitment. To determine whether residues in DnaA^DI^ are required for *oriC* opening, a DNA strand separation assay was employed {Pelliciari, 2021 #3163}. Here recombinant DnaA proteins are incubated with fluorescently labelled oligonucleotide scaffolds that mimic *oriC*, and DNA strand separation is detected using a platereader. As with experiments using a truncated DnaA protein from *Aquifex aeolicus* (17), *B. subtilis* DnaA lacking domains I and II (DnaA^III/IV^) retains strand separation activity (Fig. S10). This result supports the proposal that the essential function of these DnaA^DI^-residues is DnaD recruitment. Intriguingly, the surface of DnaA^DI^ interacting with DnaD is also the binding site for the developmentally expressed DNA replication inhibitor SirA, raising the possibility that SirA could directly compete with DnaD for binding DnaA (Fig. 6A) (27, 28).

**Figure 6.**
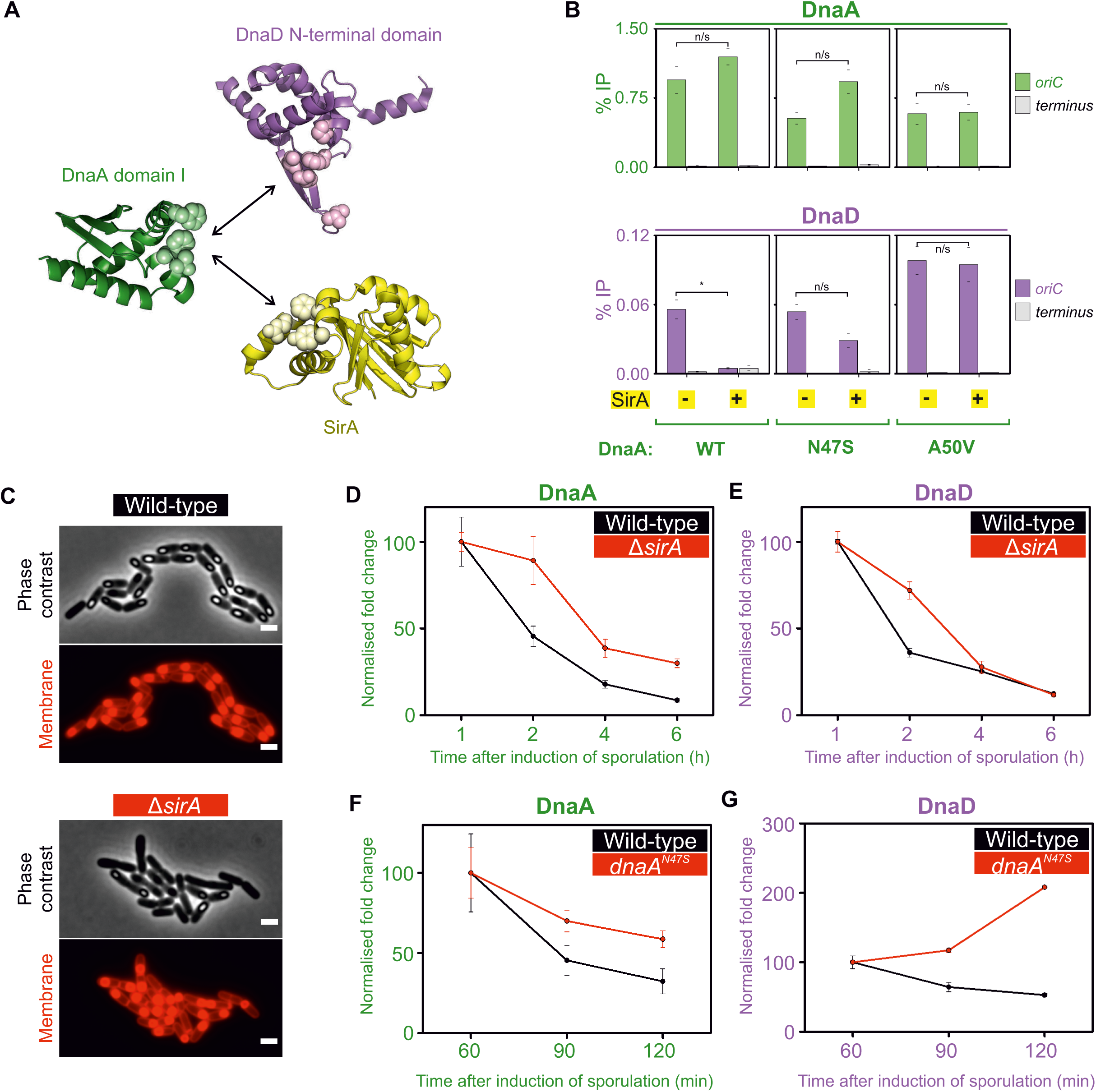
SirA binds to DnaA^DI^ and inhibits DnaD recruitment to *oriC* during sporulation. **(A)** Crystal structures of DnaD^NTD^ (PDB 2v79), DnaA^DI^ and SirA (PDB 4tps) highlighting residues at the protein:protein interfaces responsible for the DnaA^DI^-DnaD^NTD^ and DnaA^DI^-SirA interactions. **(B)** ChIP analysis showing that DnaA protein variants (wild-type (HM1565) and suppressor alleles of the DnaA-SirA interaction N47S and A50V, SF27 and SF24 respectively) remain enriched at *oriC* following overexpression of SirA, whereas DnaD is only recruited to the origin when *sirA* is not overexpressed or when using *dnaA* alleles suppressing the interaction between DnaA and SirA. * shows a p-value < 0.001 and n/s a non-significant difference. **(C)** Representative microscopy images showing that *B. subtilis* 168CA (wild-type) or cells lacking *sirA* (CW1065) were both able to sporulate by six hours after the induction of sporulation by the resuspension method. Phase bright entities show the formation of prespores in phase contrast images. Membrane staining via the Nile red dye highlights the location of spore formation within mother cells (bright red signal). **(D-E)** Depletion of DnaA **(D)** and DnaD **(E)** from the origin of replication *oriC* throughout the course of sporulation in wild-type (*B. subtilis* 168CA) or mutant cells lacking a copy of *sirA*. Fold enrichment of DnaA/DnaD at the origin compared to levels of protein recruited to the terminus was calculated as an *Ori/Ter* ratio and normalised to the 1h time point after the induction of sporulation by the resuspension method. Strains are the same as used in (C). **(F)** Depletion of DnaA from *oriC* during the early stages of sporulation in wild-type (*B. subtilis* 168CA) or mutant cells where DnaA cannot interact with SirA (*dnaA^N47S^* - Cw1073). The normalised fold change indicates that DnaA remains relatively enriched at the origin by two hours after the induction of sporulation by the resuspension method. **(G)** Enrichment of DnaD at *oriC* during the early stages of sporulation in mutant cells where DnaA cannot interact with SirA (*dnaA^N47S^* - CW1073) or depletion as expected in wild-type cells (*B. subtilis* 168CA). The normalised fold change indicates that DnaD relatively accumulates at the origin by two hours after the induction of sporulation by the resuspension method. Error bars shown in (D-G) represent propagated standard errors of the mean over three biological repeats.

### SirA binding to DnaA inhibits recruitment of DnaD and DnaB to oriC

During endospore development *B. subtilis* requires two chromosomes, one for each differentiated cell type (80). To help ensure diploidy after executing the commitment to sporulate, cells express the negative regulator of DNA replication initiation SirA (8, 81). It was proposed that SirA represses DNA replication initiation by inhibiting DnaA binding to *oriC* (28). However, SirA binds to DnaA^DI^ (27), distal to the established DNA binding motifs of DnaA. Thus, the mechanism for how SirA inhibits DnaA recruitment at *oriC* was unclear. Considering the data presented above, we explored the alternative hypothesis that SirA binding to DnaA^DI^ occludes DnaD, thereby repressing DNA replication initiation by specifically inhibiting recruitment of DnaD to *oriC*.

Using a strain containing an ectopic copy of *sirA* under the control of an IPTG-inducible promoter, SirA was expressed for 30 minutes during mid-exponential growth. Previous studies showed that DnaA-dependent replication initiation was blocked under these conditions (8, 81). ChIP of wild-type DnaA revealed stable enrichment at *oriC* following SirA expression (Fig. 6B), indicating that under these conditions SirA does not inhibit DnaA binding to DNA. ChIP of DnaD showed enrichment was abolished, consistent with the model that SirA inhibits DnaD binding to DnaA. Furthermore, enrichment of the helicase loader DnaB at *oriC*, which requires prior binding of DnaD (32), was also lost following induction of *sirA* (Fig. S11). ChIP experiments were then performed using strains harbouring the variants DnaA^N47S^ and DnaA^A50V^, which are resistant to SirA inhibition (28). Enrichment of DnaD at *oriC* was restored by DnaA^N47S^ and DnaA^A50V^, to a degree that correlated with the penetrance of the *dnaA* allele (Fig. 6B) (27). Therefore, expression of SirA during exponential growth appears to specifically inhibit DnaD recruitment to *oriC*.

### Rapid loss of DnaD enrichment at oriC during sporulation depends on SirA

To investigate the impact of SirA during sporulation, we probed the recruitment of DnaA and DnaD to *oriC* in cells induced to sporulate by the resuspension method (Fig. 6C) (82, 83). ChIP showed that in a wild-type strain, both DnaA and DnaD are depleted from *oriC* through the course of sporulation (Figs. 6D-E and S12A-B). In a Δ*sirA* mutant both proteins remained relatively enriched at *oriC* during the first two hours of the developmental process, followed by their eventual depletion (Fig. 6D-E). The apparent timing of SirA activity is consistent with previous reports indicating that *sirA* is induced early during sporulation (7, 8). Strikingly, during the first two hours of sporulation in a strain expressing the DnaA^N47S^ variant that is resistant to SirA, the relative enrichment of DnaD at *oriC* increased while for DnaA it decreased (Figs. 6F-G; S12C-D). Together, the results indicate that SirA is required for the rapid depletion of DnaD from the replication origin during the early stages of sporulation.

### SirA and DnaD compete for binding to DnaA^DI^

To investigate whether SirA and DnaD binding to DnaA^DI^ is mutually exclusive, we set up a competition experiment. A strain was engineered with ectopic copies of *sirA* and *dnaD* under the control of distinct inducible promoters (IPTG and xylose, respectively). While expression of SirA alone inhibited growth, co-expression with DnaD significantly ameliorated this effect (Figs. 7A and S13A-C). In contrast, expression of DnaD variants defective for binding DnaA (DnaD^F51A^, DnaD^I83A^ or DnaD^E95A^) could not alleviate the SirA expression phenotype (Fig. S13D-E). Marker frequency analysis confirmed that SirA expression inhibited DNA replication initiation, while co-expression with DnaD relieved this repression (Fig. 7B). Taken together, our results suggest that SirA binds to DnaA^DI^ and occludes the essential DnaA:DnaD interface (Figs. 7C), thus providing a molecular mechanism for SirA mediated inhibition of DNA replication initiation.

**Figure 7.**
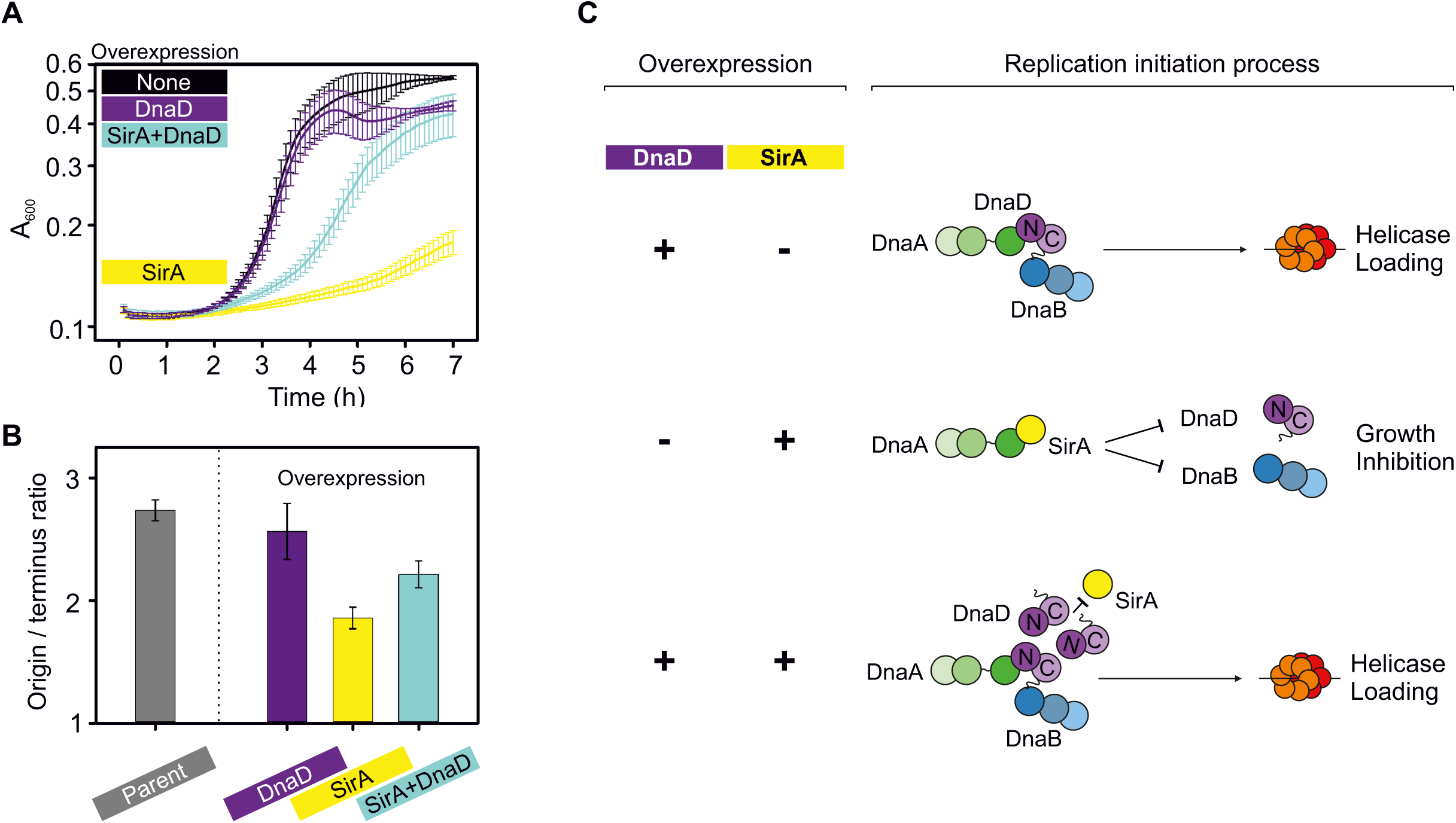
SirA and DnaD competing for binding to DnaA^DI^. **(A)** Plate reader growth assay showing that overexpression of SirA inhibits growth, which can be partially rescued by simultaneous overexpression of DnaD (CW252). None indicates cells growing without inducer, DnaD overexpression was performed with 0.35% xylose, SirA overexpression with 0.035 mM IPTG, DnaD and SirA simultaneous overexpression with 0.35% xylose and 0.035 mM IPTG. Error bars indicate the standard error of the mean for at least 3 biological replicates. **(B)** Marker frequency analysis by quantitative PCR showing that overexpression of DnaD yields a similar *ori:Ter* ratio to the control strain (168CA), whereas overexpressing SirA strongly inhibits chromosome replication and simultaneous expression with DnaD partially restores it (CW252). **(C)** Schematic outcomes of the DnaD/SirA competition assay. The sole overexpression of DnaD does not affect bacterial growth and leads to helicase loading, whereas SirA overexpression inhibits growth via binding to DnaA^DI^. This inhibition can be partially rescued and helicase loading restored when DnaD competed with SirA for DnaA^DI^ binding.

## DISCUSSION

The molecular basis underpinning how bacteria orchestrate and regulate the onset of DNA replication throughout growth and development was unclear. Here, characterization of the essential DNA replication initiation protein DnaD identified amino acids required for forming a homo-tetramer, as well as for interacting with the *B. subtilis* replication initiation proteins DnaA and DnaB (Fig. 8A). These findings illuminate specific activities required during DNA replication initiation in *B. subtilis*, which in turn engendered a model for the molecular mechanism used to regulate DNA synthesis during sporulation.

**Figure 8.**
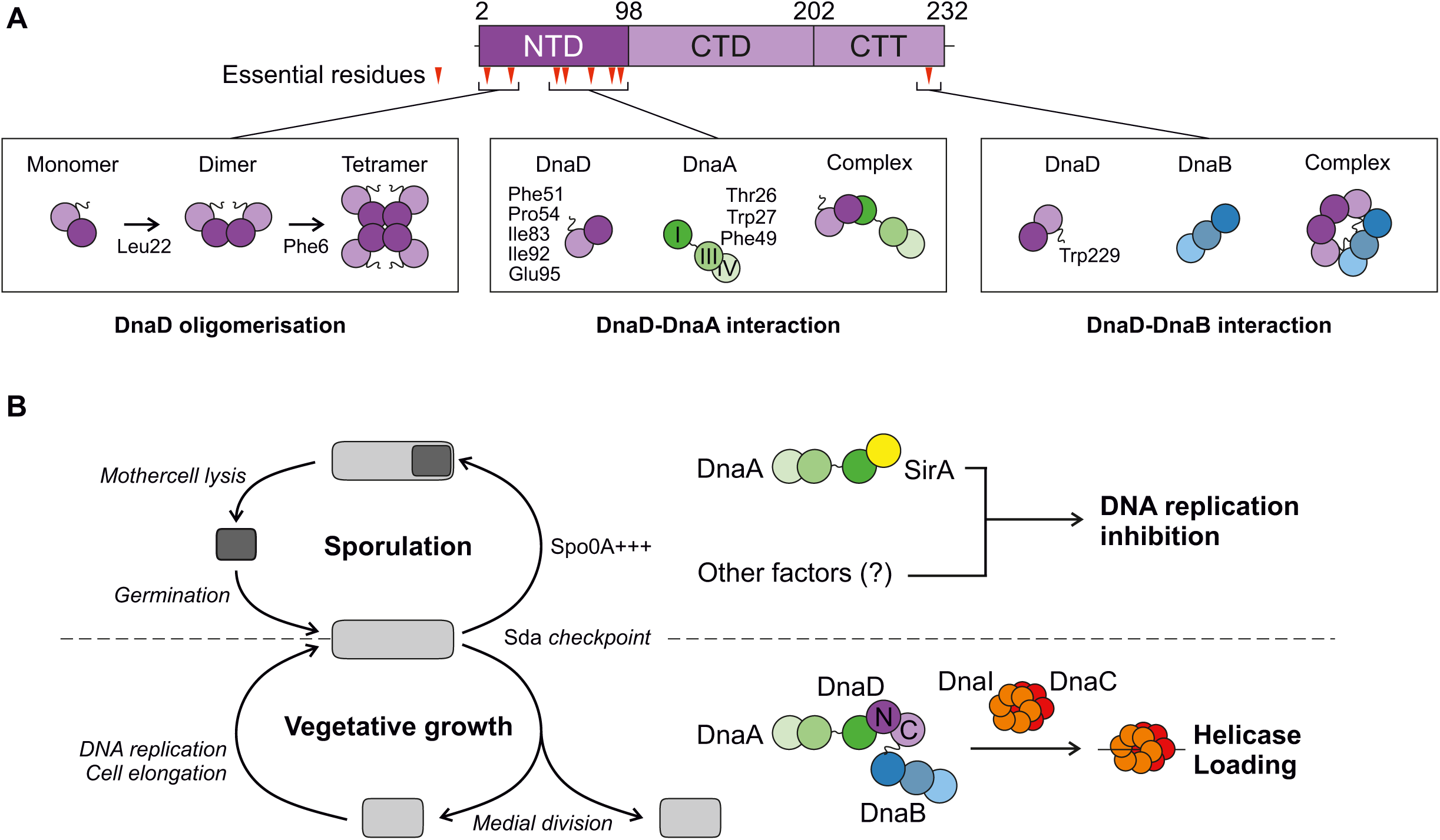
DnaD activities and interactions during DNA replication initiation at *oriC* culminate with it binding the DRE. **(A)** The DnaD functional analysis identified key residues that regulate DnaD oligomerisation and interactions with DnaA and DnaB. Essential residues are mapped as red triangles onto DnaD primary structure schematics with amino acid boundaries indicated. NTD denotes the N- Terminal Domain, CTD the C-Terminal Domain and CTT the C-Terminal Tail of DnaD. DnaD residues F6 and L22 are involved in oligomerisation, F51/P54/I83/I92/E95 in the DnaD-DnaA interaction and W229 in the DnaD-DnaB interaction. DnaA residues T26/W27/F49 are involved in the DnaA-DnaD interaction. **(B)** Proposed model for the regulation of helicase loading by SirA during *B. subtilis* sporulation or by DnaD during vegetative growth. During sporulation, DnaA can interact with SirA, possibly other unknown factors, and is gradually depleted from the origin of replication, which leads to growth inhibition. During vegetative growth, DnaA interacts with DnaD, which in turns recruits DnaB and leads to loading of the helicase complex DnaI-DnaC; this promotes replication initiation and cell growth.

The essential Leu22 was found to be specifically required for DnaD dimerization. This is in agreement with the location of Leu22 within the crystal structure of DnaD^NTD^(41). Leu22 is one of thirteen residues which are thought to become significantly buried upon dimer formation {Schneider, 2008 #2840}. We also note that while Ile17 was not essential, the DnaD^I17A^ variant did produce a growth defect (Fig. S5) and this amino acid also becomes buried upon dimerization (41).

The essential Phe6 was found to be specifically required for DnaD tetramerization. Phe6 was previously postulated to play a role in dimer:dimer formation of DnaD^NTD^ by interacting with residues in the winged-helix fold (41). Consistent with this, strains expressing alanine substitutions at Trp103 and Tyr107 (in the winged-helix) both produced growth defects (Fig. S5). Unexpectedly however, alanine substitutions at neither Thr14 nor Thr16 produced any phenotype in our screen; these had been suggested to pack onto a tight core of eight threonine side chains to stabilize the DnaD^NTD^ tetramer (41). While a molecular understanding of DnaD tetramerization requires further structural work, the identification of specific DnaD variants blocked in oligomerization that are also defective for DNA replication initiation supports the model that the functional conformation of DnaD within the cell is a tetramer.

The C-terminal tail of DnaD was found to be required for orchestrating the DnaD:DnaB interaction, particularly the essential residue Trp229 (Figs. 4A-E and 8A). Previously it was suggested that DnaB interacted with the DnaD^NTD^, although a specific binding site was not delineated (31). Based on the observation that the DnaD^CTT^ is essential and that it is required to interact with DnaB, we proposed that the DnaD^CTT^ is the critical binding site for DnaB within the cell. Furthermore, the results of two-hybrid experiments suggest that the DnaD dimer formation is required to bind DnaB, since the DnaD^L22A^ monomer did not interact with DnaB while the DnaD^F6A^ dimer did (Fig. 4E). The next step in characterizing the DnaD:DnaB interaction will be to identify the complementary binding site for DnaD on DnaB, to further our understanding of the multivalent interactions between *B. subtilis* DNA replication initiation proteins.

An extensive DnaD^NTD^ interface (Phe51, Pro54, Ile83, Ile92, Glu95; Figs. 4G-I) is essential for viability and specifically required for interacting with DnaA (Fig. 8A), consistent with two-hybrid analysis (31). The complementary interface for DnaD on DnaA^DI^ (Thr26, Trp27, Phe49) is also essential for viability, as well as for enrichment of DnaD at *oriC* (Fig. 5). Taken together, we propose that the essential role of the DnaA^DI^-DnaD^NTD^ interaction is to recruit DnaD to the replication origin.

While the DnaD alanine scan successfully pinpointed multiple protein:protein interactions, it was noteworthy that residues required for DNA binding were not isolated. Previous studies established that the DnaD^CTD^/DnaD^CTT^ contribute to DNA binding {Carneiro, 2006 #2910}{Huang, 2016 #3156}, as well as modifying the writhe of the DNA double helix (86, 87). Thus, an understanding of the role DNA binding plays for DnaD during replication initiation remains unclear.

Regulation of DNA synthesis in bacteria is mainly achieved at the stage of initiation (9). Understanding the molecular mechanisms of DNA replication initiation allows an appreciation for how regulatory factors might modulate the pathway. This was the case here for the developmental regulator SirA, where it was found to target the essential DnaA^DI^:DnaD^NTD^ interface to inhibit further rounds of DNA synthesis (Fig. 7). The SirA regulatory mechanism appears crucial to achieve rapid depletion of DnaD from *oriC* during the onset of sporulation (Fig. 6D-G). There appears to be a logic for *B. subtilis* regulating helicase loading at *oriC* during sporulation via SirA, as this mechanism would not perturb the interaction of DnaD with the replication restart primosome (85) required to ensure completion of genome replication (Fig. S14). The observation that DnaD and DnaA are eventually depleted from the chromosome origin in the absence of SirA indicates that other inhibitory systems are also present during the onset of sporulation, consistent with previous studies (84).

Lastly, homologs of *dnaD* are present in the majority of *Firmicutes,* including several clinically relevant human pathogens such as *Staphylococcus, Streptococcus, Enterococcus,* and *Listeria* (36), and conservation of essential residues in DnaD homologs suggests that these systems are analogous (Fig. S15). Moreover, replication of *S. aureus* multiresistant plasmids have been shown to require an initiation protein with structural homology to DnaD (88). The multiple essential activities of DnaD, combined with the appreciation that helicase recruitment and loading mechanisms in bacteria and eukaryotes are distinct, indicates that DnaD homologs are attractive targets for antibacterial drug development.

## DATA AVAILABILITY

All plasmids and strains are available upon request. Microscopy data reported in this paper will be shared upon request.

## ACKNOWLEDGEMENTS

SEC-MALS experiments were performed by Dr Andrew Leech at the Molecular Interaction laboratory as a service from the University of York.

## AUTHOR CONTRIBUTIONS

CW, DS, SF, SP, EM, PS, AI, HM contributed to the conception/design of the work. CW, DS, SF, SP, HM generated results presented in the manuscript. CW, DS, SF, SP created Figures. CW and HM wrote the manuscript. CW, HM, DS, SP, PS, AI edited the manuscript.

---

**Figure S1.**
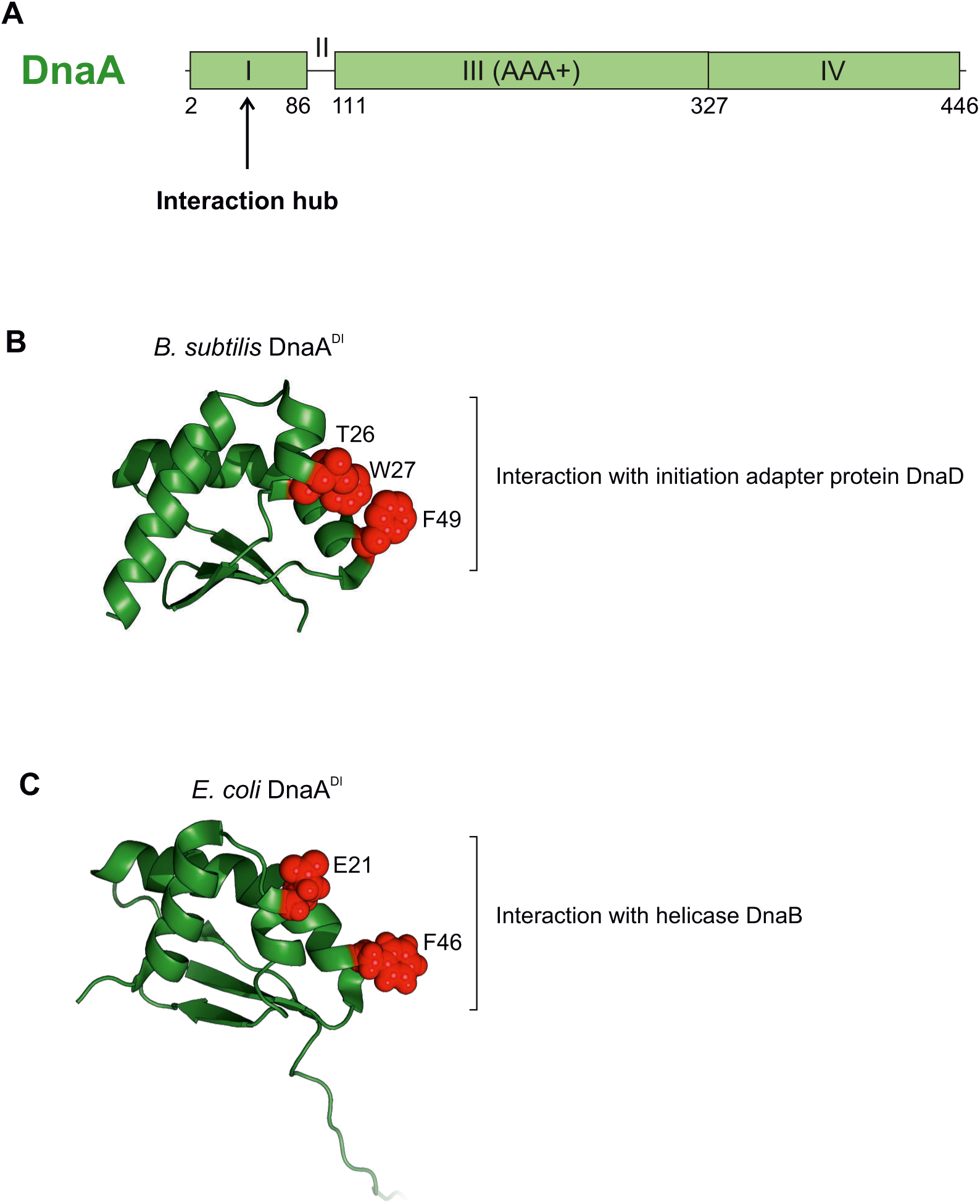
Domain organisation of DnaA highlighting a shared interaction hub in domain I. **(A)** *B. subtilis* DnaA domain organisation with amino acid boundaries indicated. **(B)** Crystal structure of *B. subtilis* DnaA domain I (PDB 4TPS) with residues thought to be involved in protein-protein interactions highlighted in red. **(C)** NMR structure of *E. coli* DnaA domain I (PDB 2E0G) with residues thought to be involved in protein-protein interactions highlighted in red.

**Figure S2.**
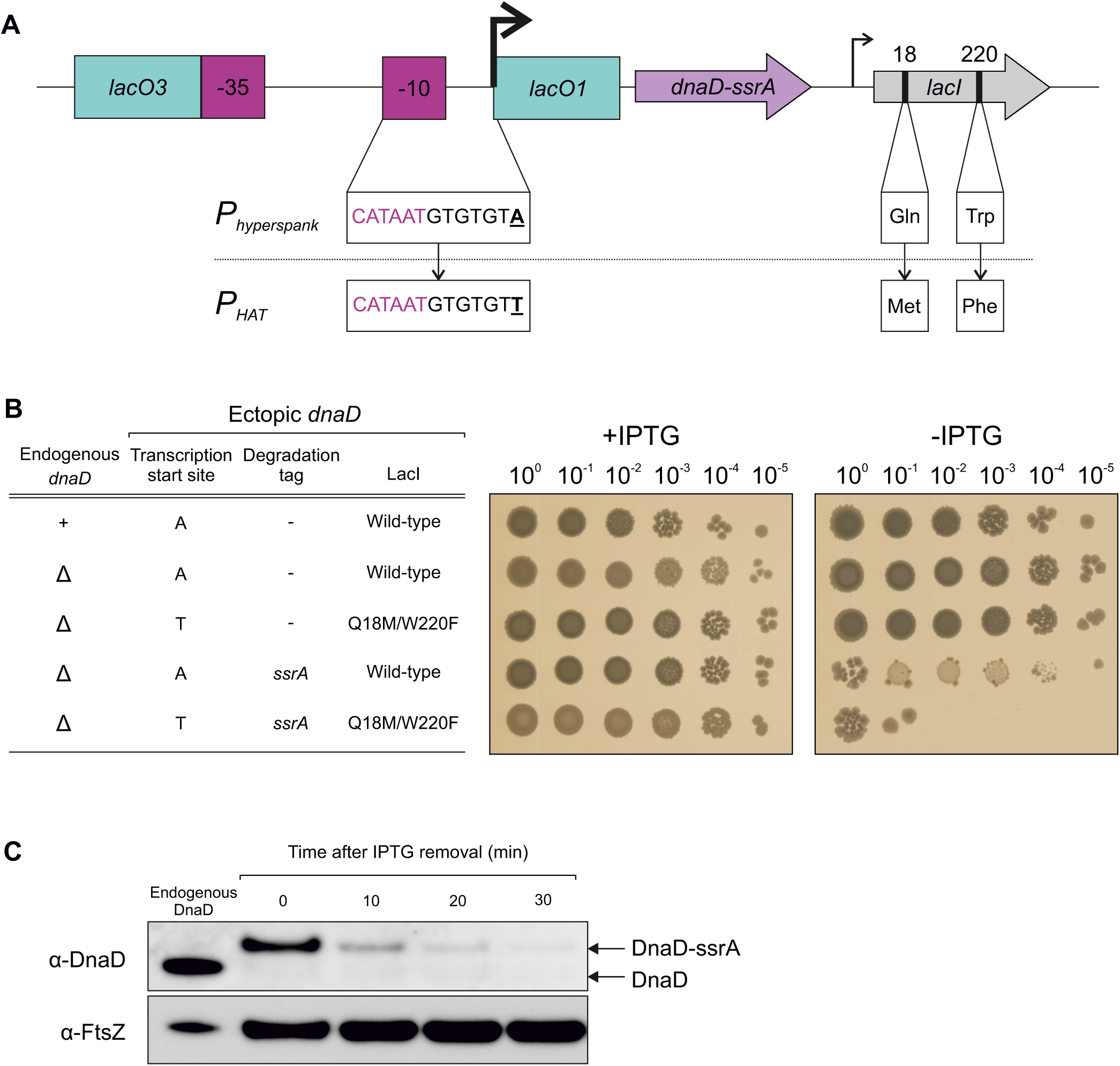
Construction of the inducible *dnaD-ssrA* strain. **(A)** Schematics of the inducible system used to drive the expression of the *dnaD-ssrA* fusion. The transcription start site (+1) of the P_hyperspank_ promoter was mutated from A to T (P_HAT_) and the *lacI* repressor binding to operator sites lacO1/lacO3 was tightened by the combination of mutations LacI^Q18M/W220F^. **(B)** Spot-titre assay showing the combination required to achieve conditional DnaD-SsrA complementation. The presence or absence of IPTG indicates that the ectopic *dnaD* copy is turned on or off, respectively. From top to bottom, strains are CW2, CW231, CW103, CW232 and CW164. **(C)** Immunoblot analysis of the inducible *dnaD-ssrA* cassette showing that degradation of DnaD-SsrA is achieved in about 30 minutes post-depletion of IPTG (CW164); endogenous *dnaD* indicates expression of wild-type DnaD in *B. subtilis* 168CA. Detection of the tubulin homolog FtsZ was used as a loading control.

**Figure S3.**
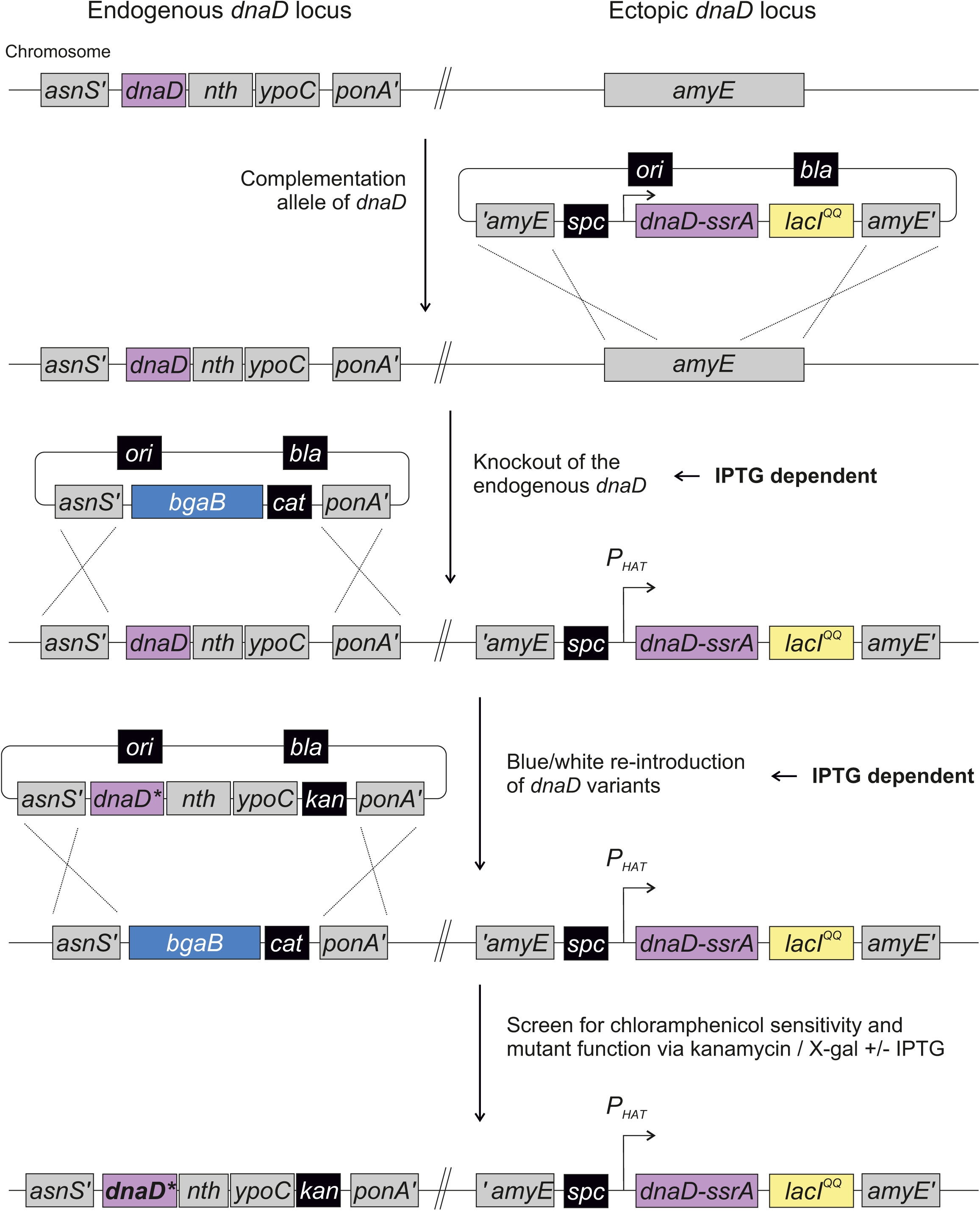
Methodology for genetic complementation and introduction of *dnaD* mutants. Schematics of the blue/white screening assay. In *B. subtilis* 168CA, the inducible complementation cassette *dnaD-ssrA* was inserted at the *amyE* locus (CW162), followed by replacement of the native *dnaD* operon by a *bgaB* cassette (encoding β-galactosidase, CW197). In the presence of the chromogenic substrate X-gal, colonies containing the *bgaB* cassette appear blue. Selection of *dnaD* mutants is performed in the presence of kanamycin, X-gal (blue/white) and IPTG (functional complementation). The function of individual mutants is then tested by growing cells in the presence or absence of IPTG.

**Figure S4.**
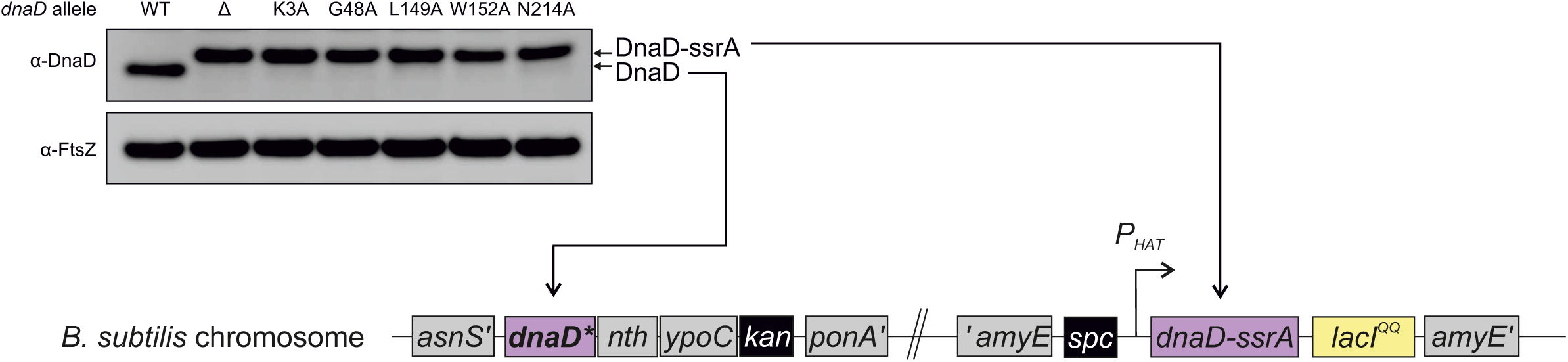
Essential DnaD residues that are not expressed *in vivo.* Immunobloting shows that some lethal alanine substitutions in the endogenous copy of DnaD were not well expressed *in vivo*. Only the expression of the ectopic DnaD-SsrA copy could be detected, whereas expression of the mutants was comparable to a strain lacking an endogenous copy of *dnaD*. Detection of the tubulin homolog FtsZ was used as a loading control. Wild-type (*B. subtilis* 168CA), Δ (CW197), K3A (CW289), G48A (CW293), L149A (CW308), W152A (CW302), N214A (CW283). Relevant schematics of the chromosome show which proteins are associated with the detected DnaD bands on the immunoblot.

**Figure S5.**
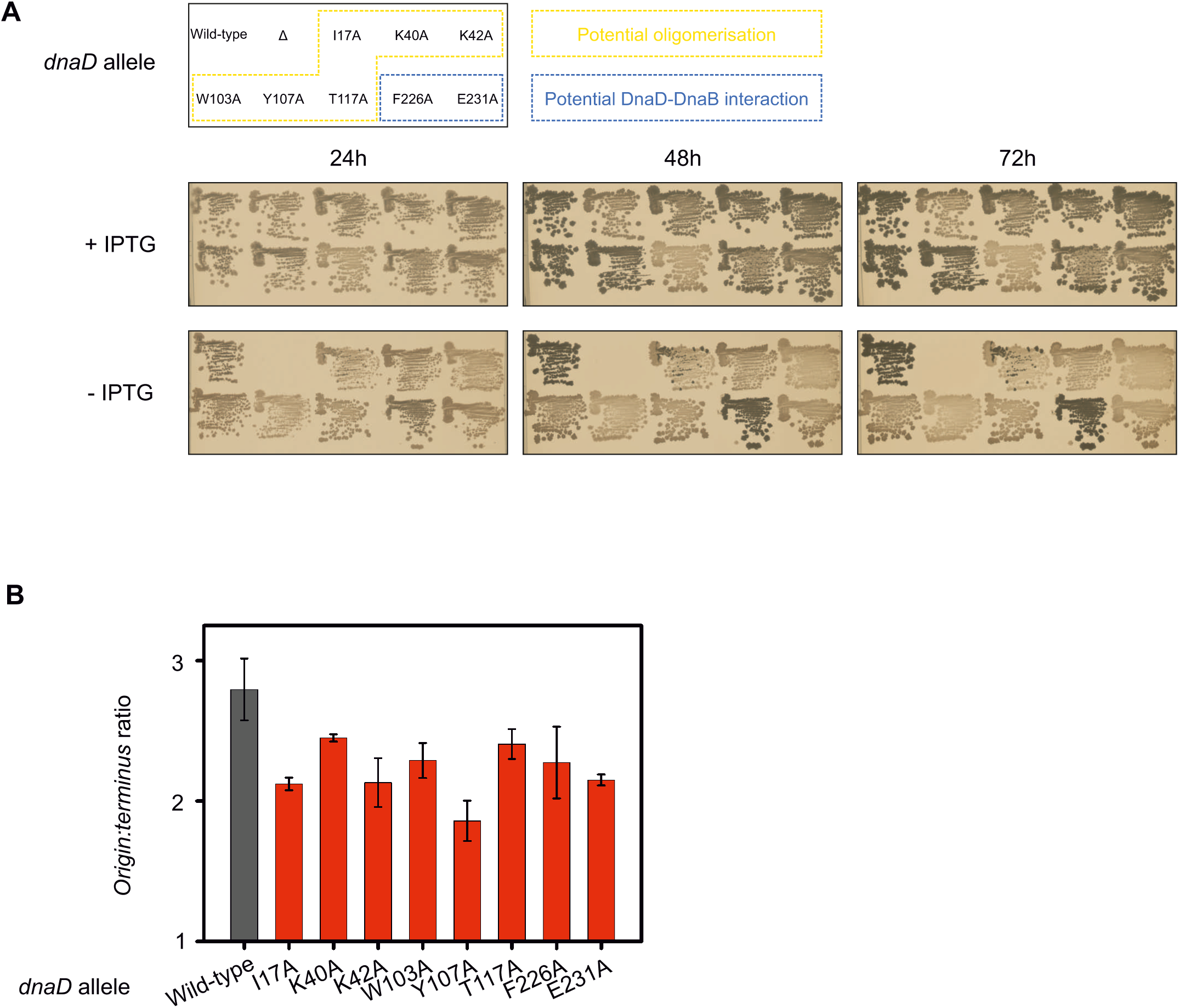
Analysis of DnaD intermediate phenotype mutants. **(A)** Colony restreaks of DnaD mutants that are associated with a translucent phenotype or leading to heterogeneous colonies over 72h growth at 37°C. Plates are shown in the presence (+IPTG) or absence (- IPTG) of *dnaD-ssrA* and their corresponding layout is detailed at the top (WT indicates wild-type). The location of these residues in DnaD structure suggests they are involved in oligomerisation or contribute to the DnaD-DnaB interaction. Wild-type (CW162), Δ (CW197), I17A (CW290), K40A (CW306), K42A (CW307), W103A (CW298), Y107A (CW299), T117A (CW300), F226A (CW284), E231A (CW309). **(B)** Marker frequency analysis by quantitative PCR associated with strains shown in panel (A) in the absence of DnaD-SsrA. Primers used to amplify the origin annealed within the *incC* region. Strains are the same as shown in (A).

**Figure S6.**
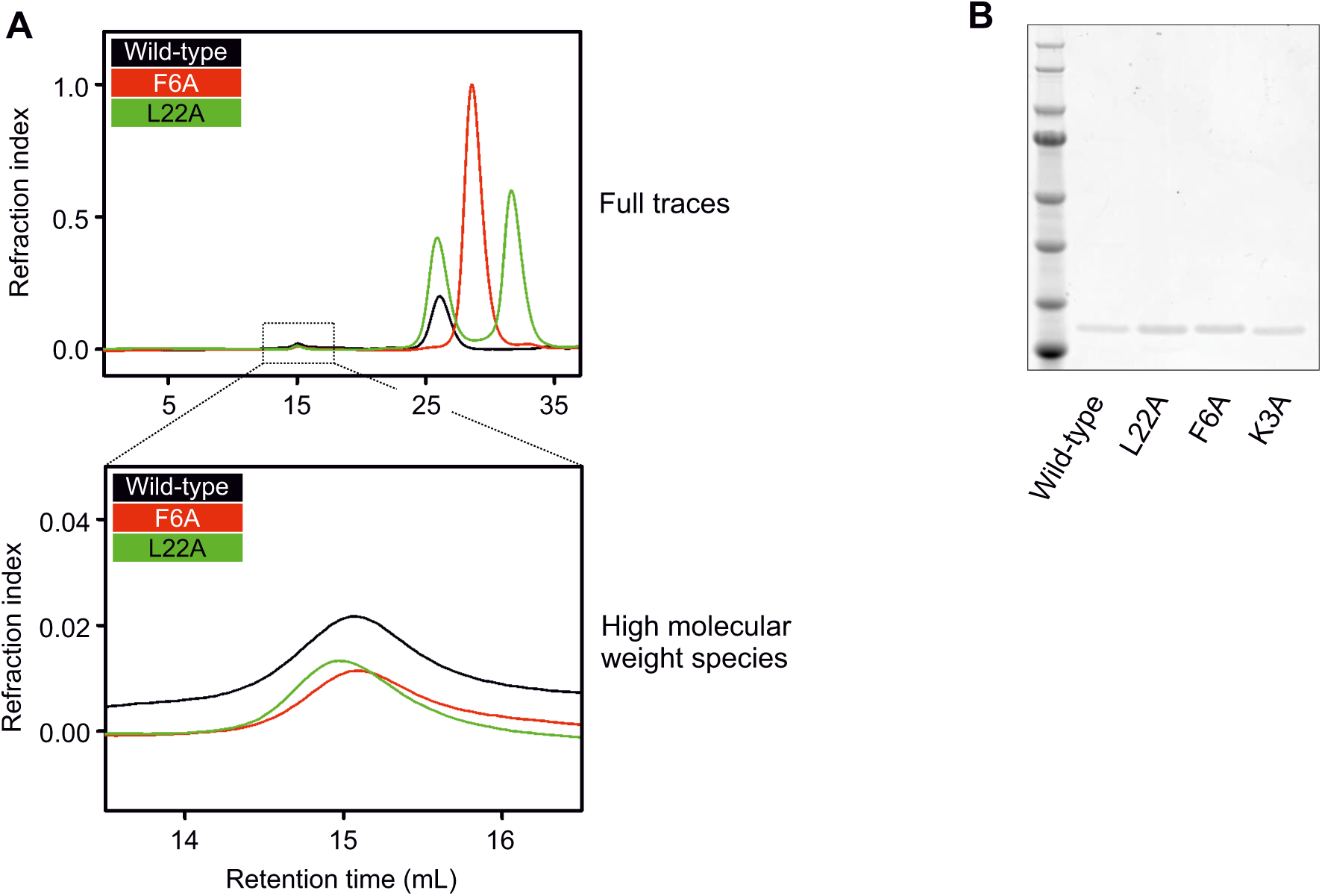
Size-exclusion chromatography and crosslinking analysis controls. **(A)** Full trace of the UV-spectrum represented as a refraction index during the SEC experiments showing that the DnaD variants (wild-type, F6A and L22A) did not display major fractions of aggregates (high molecular weight species). **(B)** Coomassie staining of the DnaD variants (wild-type, L22A, F6A and K3A) that were used during the BS3 crosslinking assay showing that all species were approximately used in equimolar amounts.

**Figure S7.**
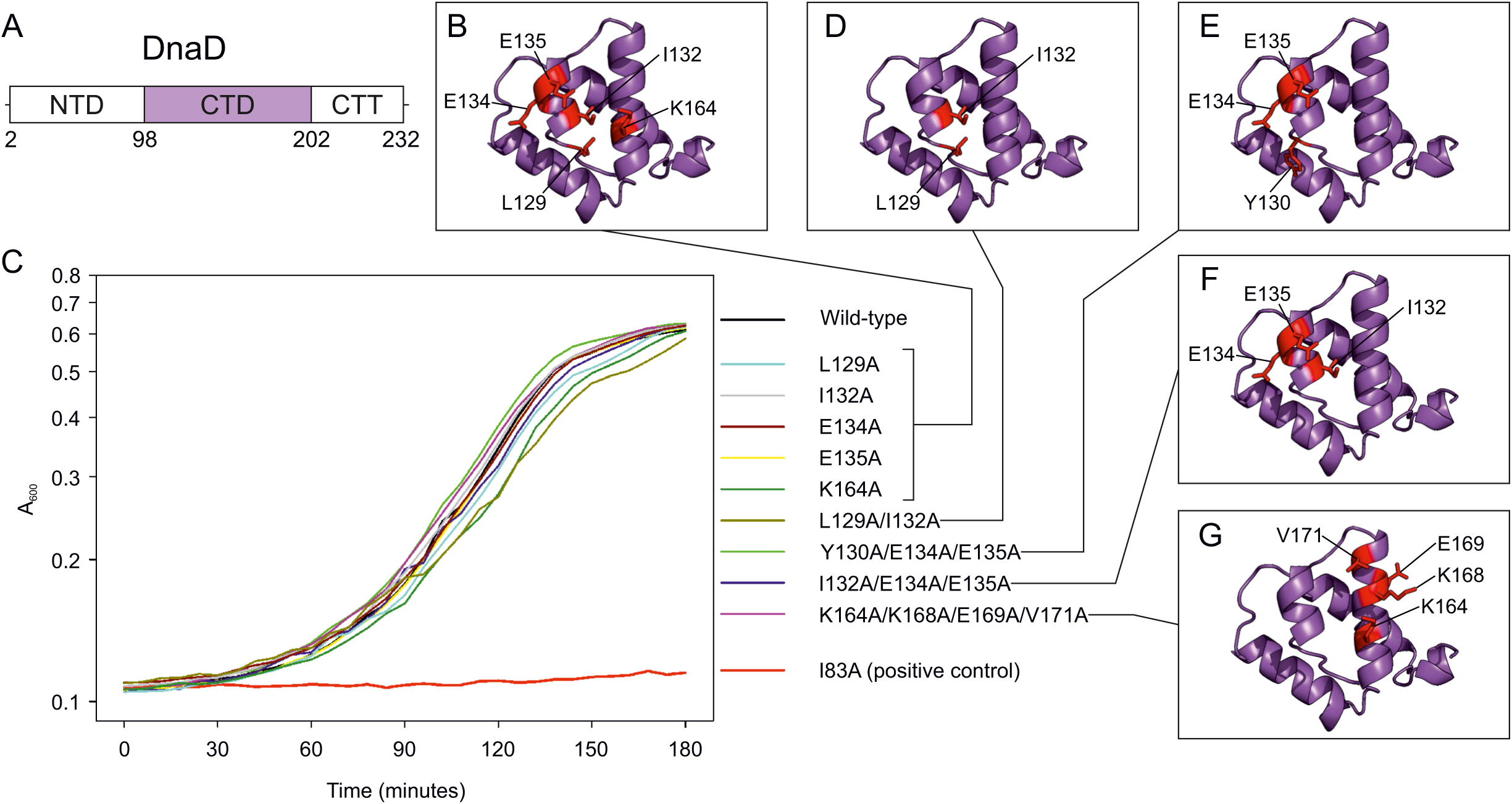
Residues in DnaD C-terminal domain that interact with DnaA^DI^ *in vitro* are not essential *in vivo*. **(A)** Domain organisation of DnaD with amino acid boundaries indicated. NTD denotes the N-Terminal Domain, CTD the C-Terminal Domain and CTT the C-Terminal Tail of DnaD. **(B)** Individual substitutions in DnaD^CTD^ mapped onto the NMR structure. **(C)** Growth analysis of *B. subtilis* DnaD variants using the inducible *dnaD-ssrA* strain in the absence of IPTG. Individual or combinations of mutations in DnaD^CTD^ show that growth was unaffected by these changes. Strains in legend (from top to bottom) are: CW162, CW179, CW167, CW171, CW172, CW173, CW176, CW177, CW178, CW168, CW170. **(D-G)** Multiple substitutions in DnaD C-terminal domain that were used in (C) mapped onto the NMR structure (37).

**Figure S8.**
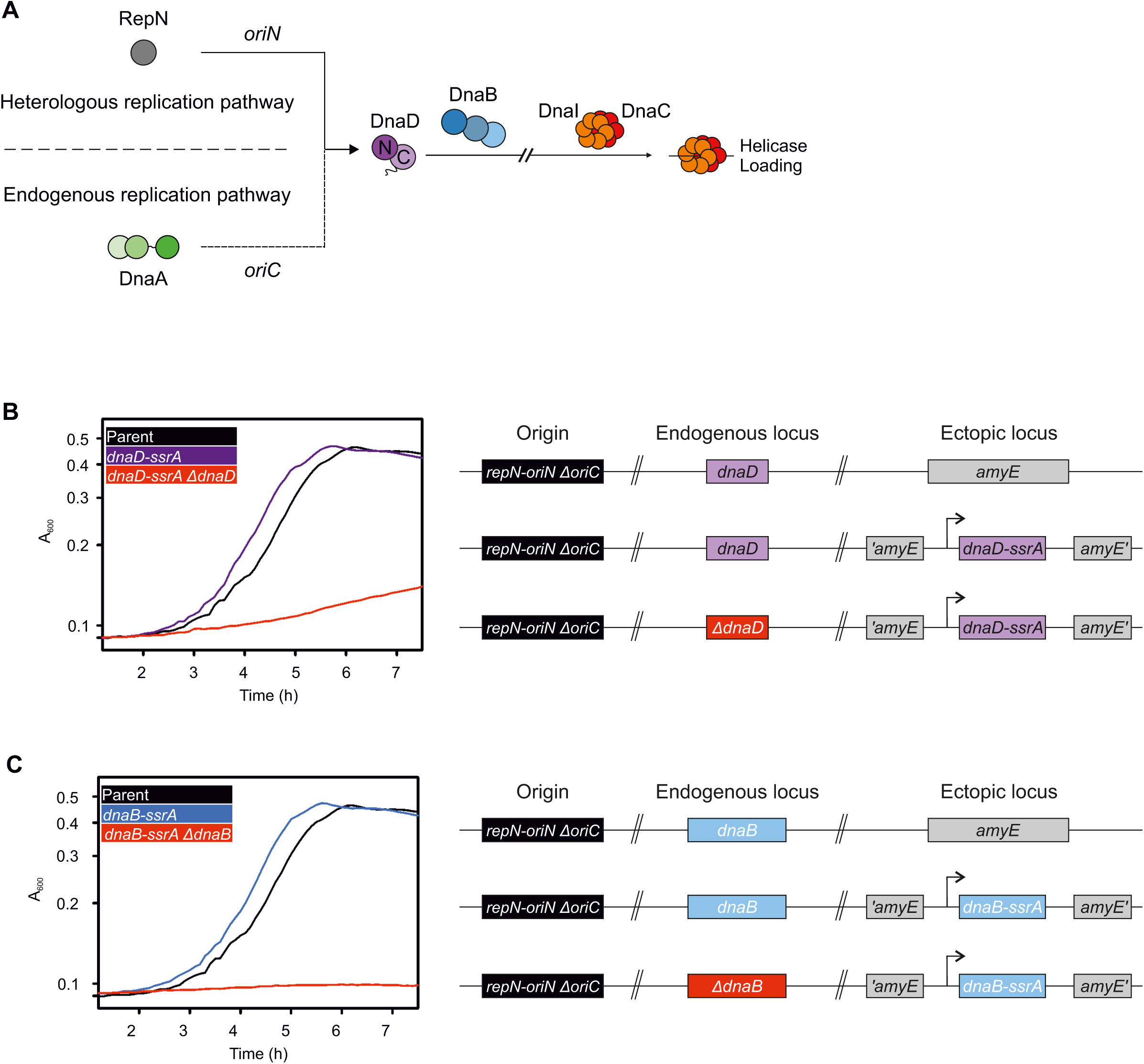
Endogenous and heterologous DNA replication systems used for the study of DnaA variants in *B. subtilis*. **(A)** Endogenous replication via DnaA at *oriC* can be complemented by the presence of the heterologous *oriN-repN* replication system. Here, this allows the study of DnaA or origin mutants without acquiring suppressor mutations. Note that both pathways require DnaD and DnaB to achieve helicase loading. **(B)** Plate reader assay showing growth of a strain replicating exclusively via *oriN,* with (*dnaD-ssrA*, CW651) and without (*dnaD-ssrA ΔdnaD*, CW658) DnaD expression. **(C)** Plate reader assay showing growth of a strain replicating exclusively via *oriN,* with (*dnaB-ssrA*, CW652) and without (*dnaB-ssrA ΔdnaB*, CW659) DnaB expression. Parent in (B-C) shows the growth profile of a strain replicating exclusively via *oriN* that only contains the native copies of *dnaD* and *dnaB* (HM950).

**Figure S9.**
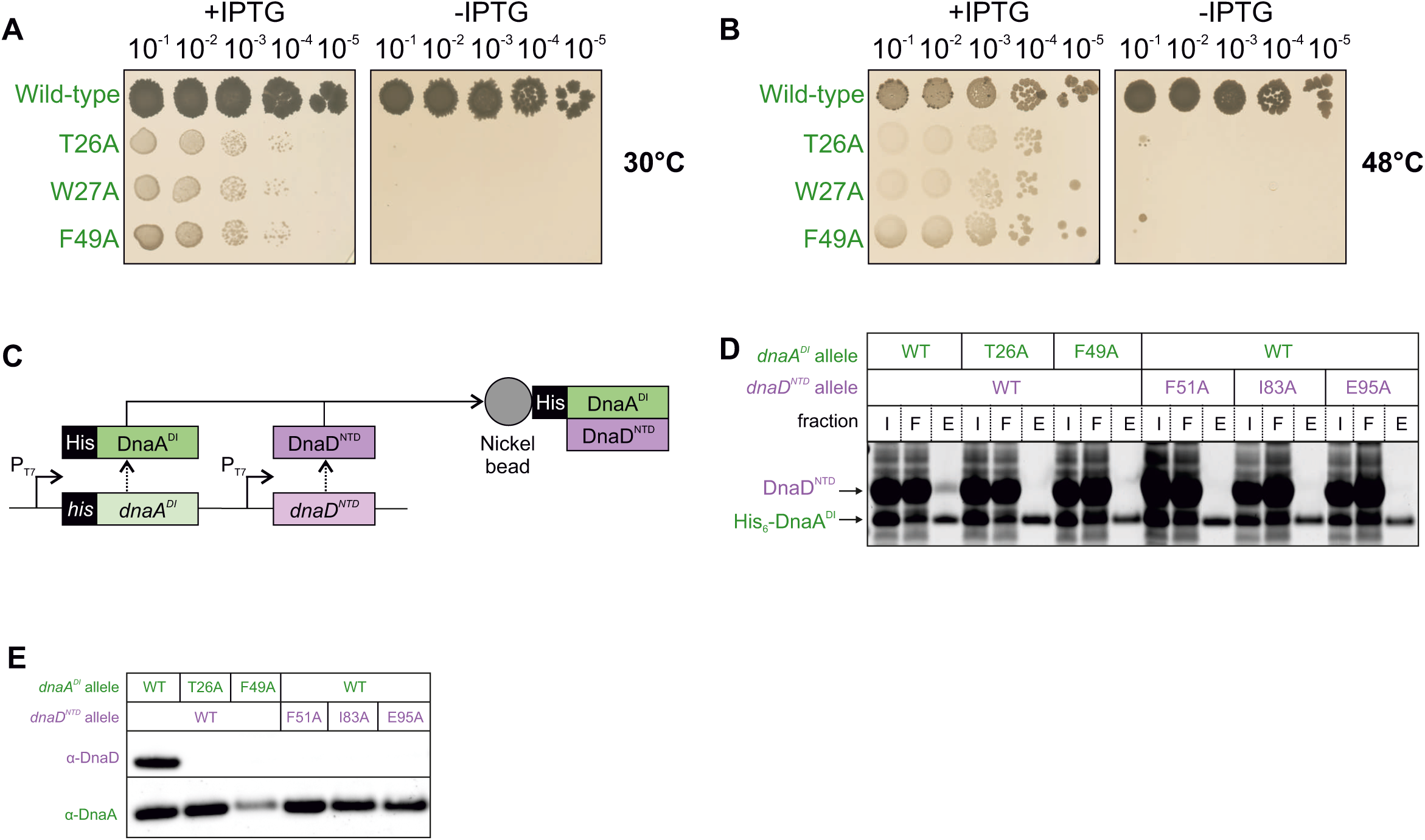
Residues in DnaA^DI^ disrupt the interaction between DnaA and DnaD. **(A-B)** Spot titre analyses showing that DnaA^DI^ variants produced a lethal phenotype *in vivo*. The presence or absence of IPTG indicates the induction state of the *repN/oriN* system. Wild-type (HM1108), T26A (HM1540), W27A (HM1541), F49A (HM1542). **(A)** shows DnaA domain I mutants lethal phenotype after 72h growth at 30°C. **(B)** shows DnaA domain I mutants lethal phenotype after 48h growth at 48°C. **(C)** Schematic of the pull-down assay using *his -dnaA^DI^* and *dnaD^NTD^* to probe for an interaction between DnaA domain I and the N-terminal domain of DnaD using Nickel beads. **(D)** Eluate staining from DnaA-DnaD pull-down assays showing loss of interaction between His -DnaA^DI^ and DnaD^NTD^ when using variants of DnaA or DnaD. WT indicates wild-type proteins and Input, Flow through and Eluate fractions are respectively indicated as I, F and E. **(E)** Immunoblot analysis of DnaA and DnaD mutant overexpression eluates from pull-down assays showing the identity of each polypeptide. Samples are the same as used in panel (D). Plasmids used in (D-E) are listed in Table S2.

**Figure S10.**
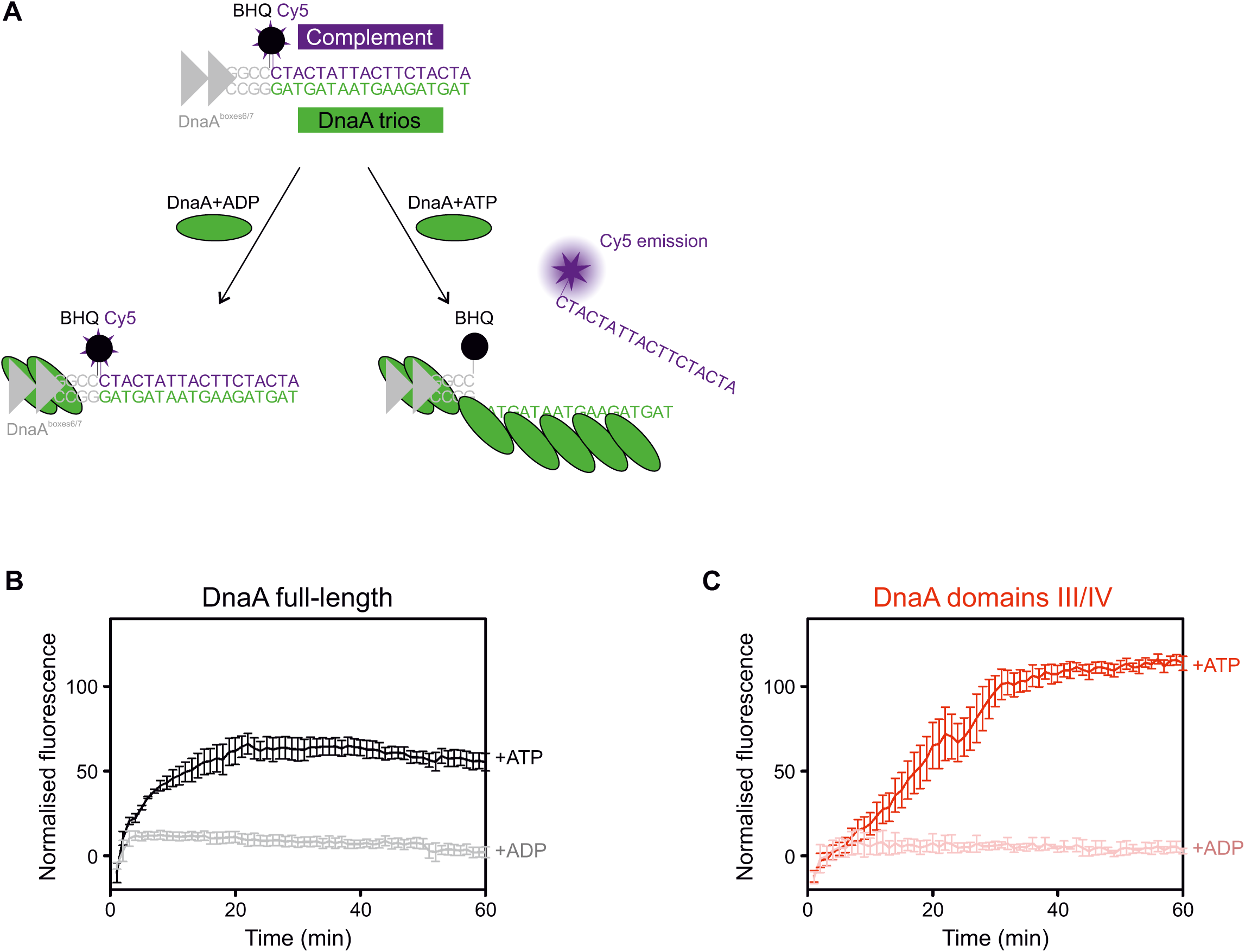
DnaA does not require domains I and II to unwind DNA substrates. **(A)** Illustration of the strand separation assay setup used to detect DnaA-directed unwinding of DNA substrates. Three oligonucleotides are annealed to mimick the *B. subtilis* origin unwinding region including DnaA boxes (dsDNA binding via DnaA) and the DnaA trios/complementary region. The bottom strand is continuous and unlabelled. The top strand corresponding to the DnaA boxes and GC-rich region is labelled with a black- hole quencher (BHQ) at the 3’-end and the complementary oligonucleotide to the trios is labelled with Cy5 at the 5’-end. As a fully dsDNA probe, the BHQ quenches fluorescence emitted by the Cy5 group. Upon incubation with DnaA and ADP, DnaA binds DnaA boxes, cannot engage the DnaA trios and no fluorescence remains quenched. In the presence of ATP, DnaA binds DnaA boxes and forms an oligomer on the DnaA trios, thereby displacing the probe complementary to the trios and allowing emission of Cy5 fluorescence. **(B-C)** Strand separation assays performed with the same probe in the presence of protein variants of DnaA. DNA substrate: oHM558/oHM778:oHM590. Background corresponding to the basal fluorescence of the DNA probe was subtracted from the curves. Error bars show the standard error of the mean for three biological replicates. **(B)** shows that DnaA full-length is able to separate strands in the presence of ATP. **(C)** shows that a truncation of DnaA lacking domains I and II retained the ability to separate strands in the presence of ATP.

**Figure S11.**
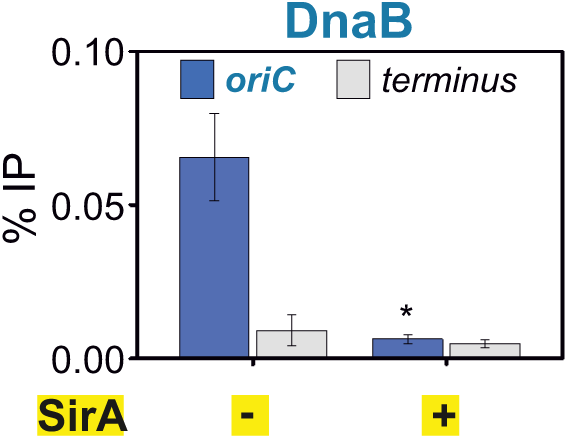
SirA overexpression abolishes DnaB recruitment to *oriC*. ChIP analysis showing that DnaB recruitment to *oriC* is lost following overexpression of SirA (HM1565). Primers used to amplify the origin annealed within the *incC* region. * shows a p-value of 0.0277.

**Figure S12.**
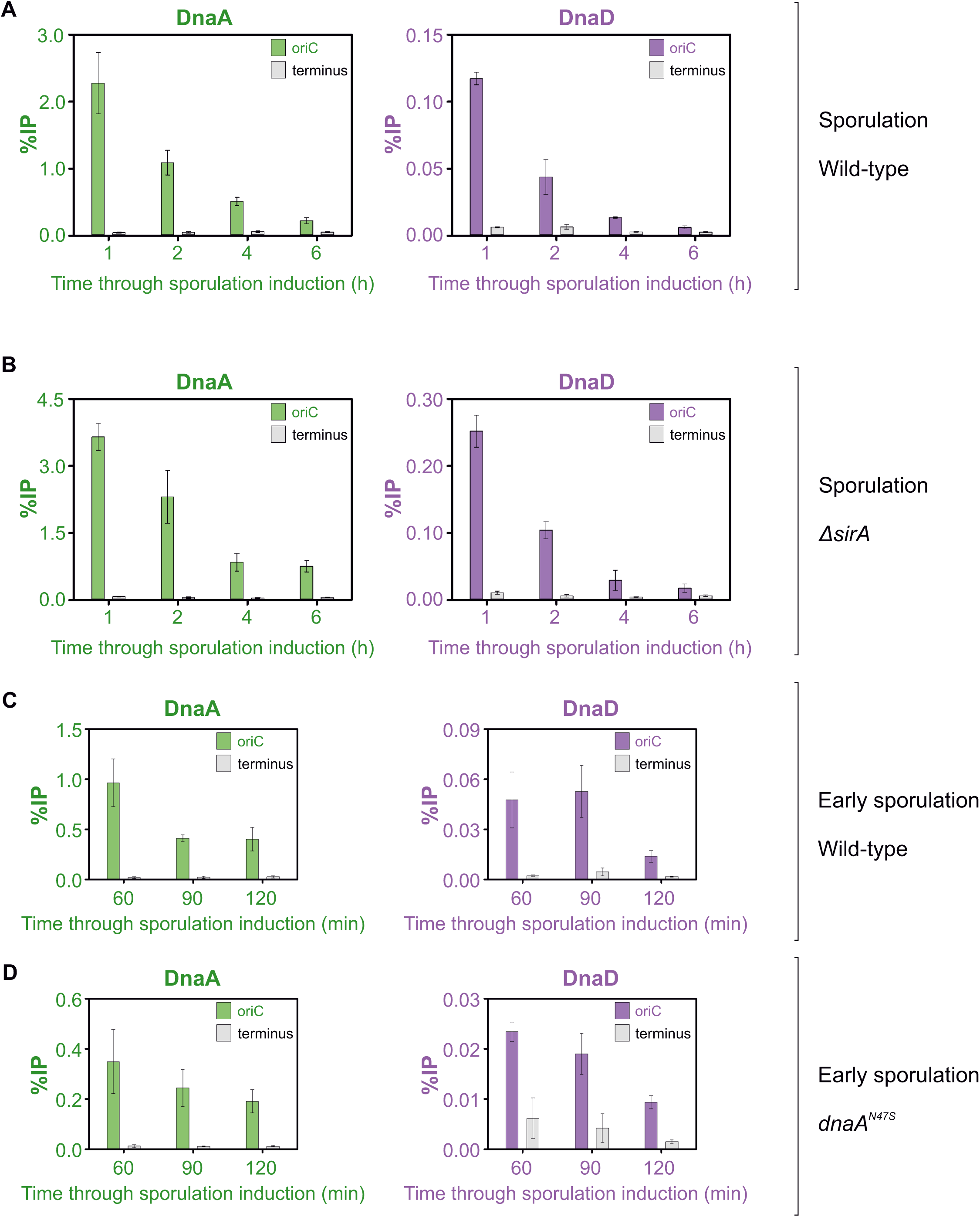
DnaA and DnaD are depleted from the origin during sporulation. **(A-B)** ChIP of DnaA and DnaD proteins at *oriC* throughout the induction of sporulation **(A)** in a wild-type strain (*B. subtilis* 168CA) and **(B)** in a knockout strain of *sirA* (CW1065). **(C-D)** ChIP of DnaA and DnaD proteins at *oriC* during early sporulation **(C)** in a wild-type strain (*B. subtilis* 168CA).and **(D)** in a strain suppressing the interaction between DnaA and SirA using the *dnaA^N47S^* allele (CW1073). Primers used to amplify the *oriC* in (A-D) annealed within the *incC* region. Error bars show the standard error of the mean over three biological repeats.

**Figure S13.**
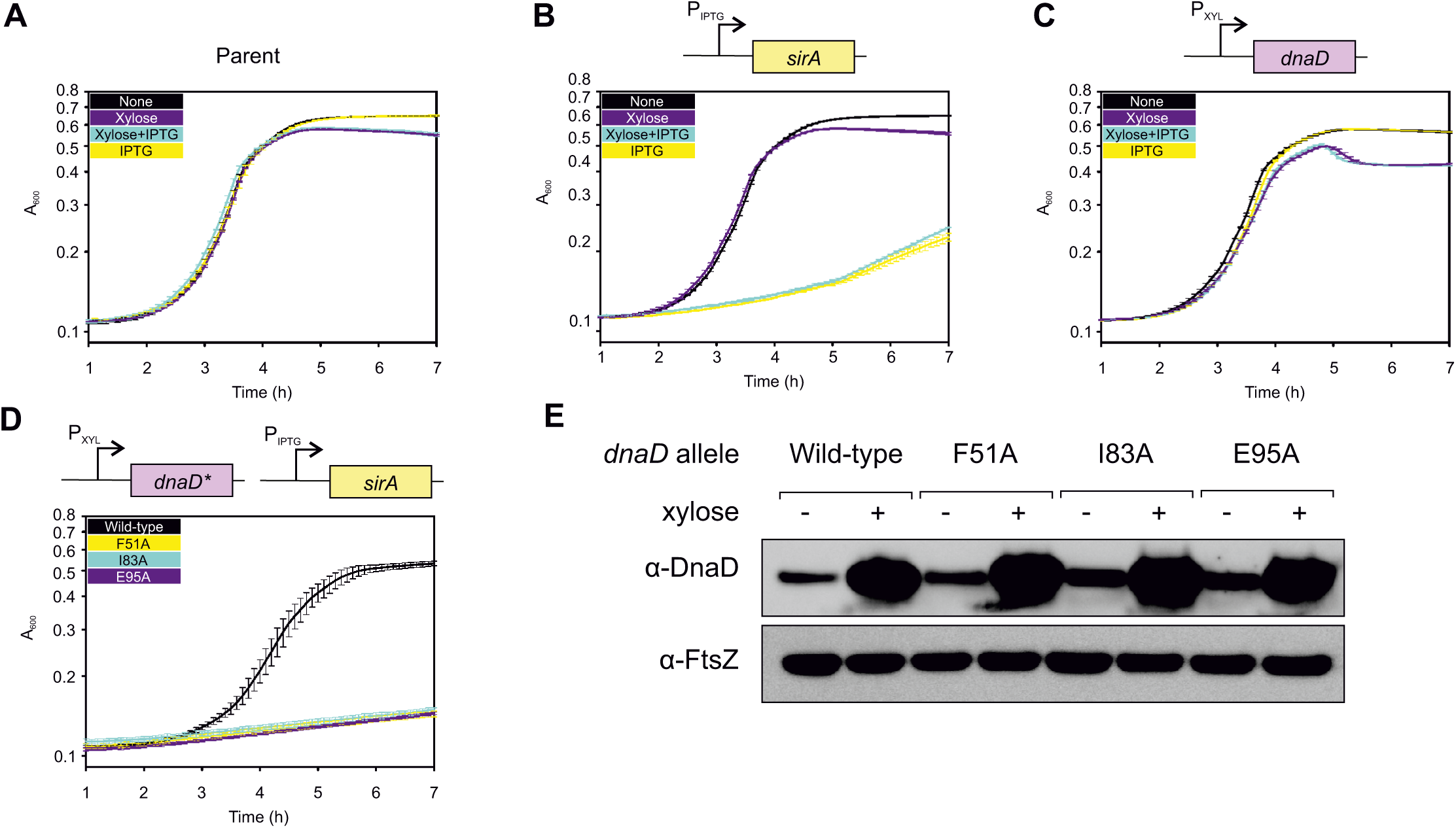
SirA inhibits the DnaA:DnaD interaction by preventing DnaD recruitment to *oriC*. **(A-C)** Plate reader growth assays with either no inducer (None), Xylose at 0.35%, IPTG at 0.035 mM or Xylose and IPTG together. **(A)** Shows that wild type *B. subtilis* 168CA grows in all conditions. **(B)** Shows that SirA overexpression in a strain background lacking the DnaD overexpression cassette inhibits bacterial growth, and that this inhibition is solely due to the addition of 0.035 mM IPTG (CW260). **(C)** Shows that DnaD overexpression in a strain background lacking the SirA overexpression cassette does not affect bacterial growth (CW261). **(D)** Plate reader analysis in the presence of xylose (0.35%) and IPTG (0.035 mM) shows that DnaD^NTD^ mutants F51A, I83A and E95A do not rescue SirA-dependent growth inhibition. Wild-type (CW252), F51A (CW279), I83A (CW270), E95A (CW280). Error bars in (A-D) indicate the standard error of the mean for two biological replicates. **(E)** Immunoblot analysis showing that DnaD variants were overexpressed to similar levels following xylose induction (0.035%). Detection of the tubulin homolog FtsZ was used as a loading control. Strains are the same as used in (D).

**Figure S14.**
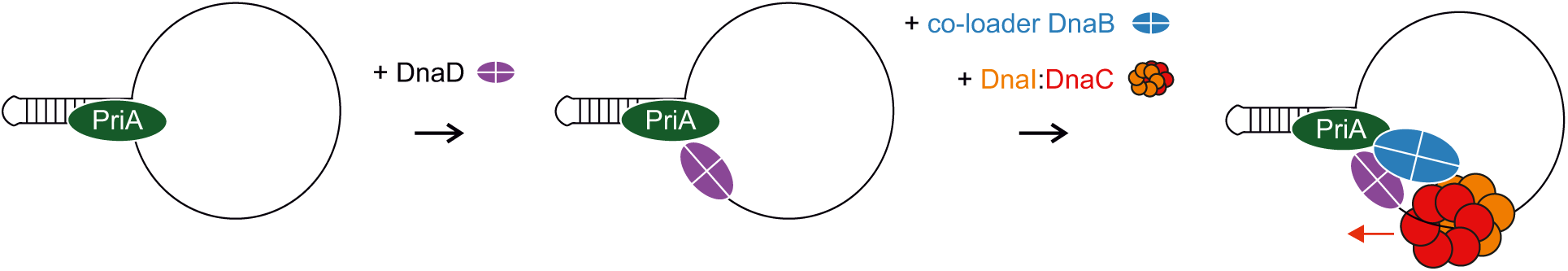
Model for helicase recruitment and loading in *B. subtilis* during PriA-dependent replication restart at a single-strand origin (*sso)*.

**Figure S15.**
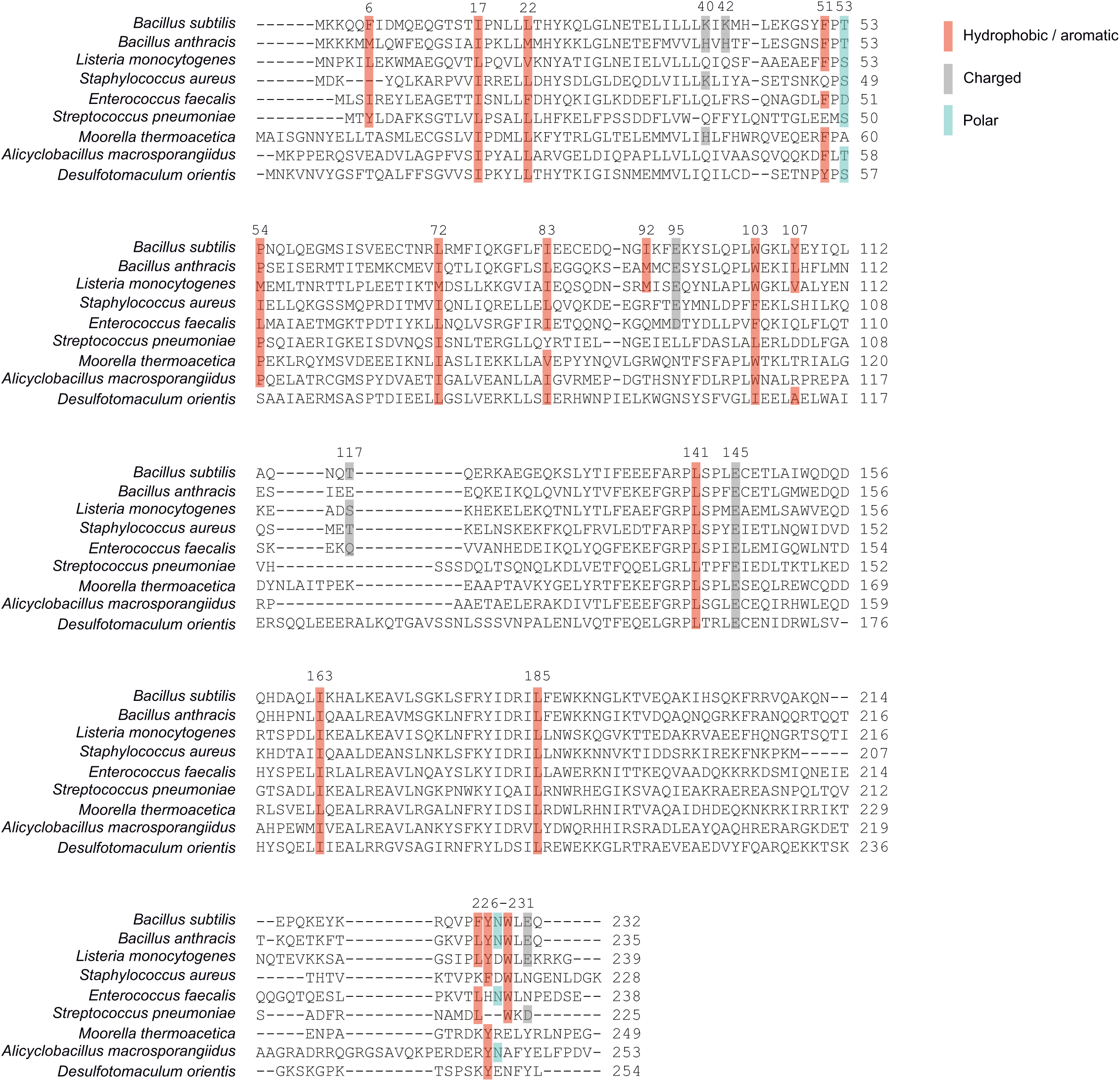
DnaD protein sequence conservation. Protein alignment showing the conservation of *B. subtilis* DnaD residues that displayed a phenotype *in vivo* and remained expressed. Numbers by the end of each row indicate amino-acid positions with respect to specific species and those on top of coloured boxes (red for hydrophobic / aromatic, grey for charged and blue for polar amino acid classes) are relative to the *B. subtilis* DnaD sequence.

